# Amacrine cells differentially balance zebrafish colour circuits in the central and peripheral retina

**DOI:** 10.1101/2022.01.22.477338

**Authors:** Xinwei Wang, Paul A Roberts, Takeshi Yoshimatsu, Leon Lagnado, Tom Baden

**Author notes:** Correspondence to (XW), (LL), (TB). Twitter: @XinweiWang1017 (XW), @PAR_EyeMath (PAR), @TaYoshimatsu (TY), @Neonsynapse (LL), @NeuroFishh (TB). Equal contribution (Leon Lagnado, Tom Baden). **Author contributions.** Conceptualization, XW, LL, TB; Methodology, XW, PAR, TY; Investigation, XW, PAR; Data Curation, XW, TB; Writing – Original Draft, TB; Writing – Review & Editing, XW, PAR, TY, LL, TB; Visualization, XW, TB; Supervision, LL, TB; Project Administration, LL, TB; Funding Acquisition, LL, TB. The authors declare no conflict of interest.

## Abstract

In vertebrate vision, the feature-extracting circuits of the inner retina are driven by photoreceptors whose outputs are already pre-processed. In zebrafish, for example, outer retinal circuits split “colour” from “greyscale” information across all four cone-photoreceptor types. How does the inner retina process this incoming spectral information while also combining cone-signals to shape new greyscale functions?

We address this question by imaging the light driven responses of amacrine cells (ACs) and bipolar cells (BCs) in larval zebrafish, in the presence and pharmacological absence of inner retinal inhibition. We find that amacrine cells exert distinct effects on greyscale processing depending on retinal region, as well as contributing to the generation of colour opponency in the central retina. However, in the peripheral retina amacrine cells enhanced opponency in some bipolar cells while at the same time suppressing pre-existing opponency in others, such that the net change in the number of colour-opponent units was essentially zero. To achieve this ‘dynamic balance’ ACs counteracted intrinsic colour opponency of BCs via the On-channel. Consistent with these observations, Off-stratifying ACs were exclusively achromatic, while all colour opponent ACs stratified in the On-sublamina.

This study reveals that the central and peripheral retina of larval zebrafish employ fundamentally distinct inhibitory circuits to control the interaction between greyscale- and colour-processing. Differential actions on the On- and Off-channels control the transmission of colour-opponent signals in the periphery.

## INTRODUCTION

Animal eyes encode patterns of light along distinct axes of variation such as space, time and “colour”^1^. These axes combine signals from a shared population of photoreceptors, which poses a general question in neural circuit organisation: how can a common set of inputs be processed such that specialisation for one task does not simultaneously deteriorate function elsewhere?

One strategy is to split the neural signal down separate microcircuits that implement different processing tasks. Such parallel processing is fundamental to brain function^2^, including in the vertebrate retina^3,4^, where stimulus-response relationships become increasingly specific and diverse as the visual signal travels from photoreceptors via bipolar cells (BCs) to retinal ganglion cells (RGCs). Based on this architecture, progressive circuit specialisation for distinct tasks can then take place in different populations of inner retinal neurons. A key question then is: at what stage in the retinal circuit does specialisation for each processing task occur? Specialisation for one task may precede specialisation for others. In teleost fish, for example, spectral coding in the outer retina^5^ precedes the extraction of key spatiotemporal features of the inner retina^6^.

In larval zebrafish, splitting of “colour” and “brightness” signals begins in the outer retina, where the cone-photoreceptors are modulated by horizontal cells at the first synapse in vision^7^. Red-cones provide an output that signals brightness and is essentially colour-invariant, while green-cones provide a primary colour output that is brightness-invariant. Further, blue- and UV-cones provide secondary colour and brightness channels, respectively. Such parallel representation^7–10^ of spectral information is beneficial for colour vision^11^, but how can it be maintained through the dense interconnectedness of inner retinal circuits^12–14^? All the various processing channels evident in the retinal output involve amacrine cells (ACs) in the inner retina that modify the visual signal by inhibition through both GABA- and glycinergic transmission alongside a variety of neuromodulators that can either inhibit or potentiate transmission^14–17^. How do these operations on the visual signal in the inner retina adjust the results of spectral processing arriving from the outer retina? Distinct types of BC do represent the spectral inputs from each of the four cone-signals in isolation, but alongside them are a plethora of other BCs that represent diverse cone-mixtures, presumably specialized for other temporal and spatial processing tasks^18,19^.

ACs are the most diverse yet least understood class of neurons in the retina^6,20,21^. In mice, transcriptomic analysis revealed 63 molecularly distinct types of ACs^22^, while in zebrafish, >20 can be defined by anatomy^12^. General roles of ACs include the shaping of BC and RGC receptive field structures^14,20^, modulating their dynamic range^23,24^ and adaptive properties to generate the functional diversity of the retinal output^13,25,26^. Although the specific functions of most AC-types across any species remains unknown, we might expect ACs to contribute to both chromatic and achromatic signalling^27–29^. To test if this is the case, we surveyed light-driven signals of ACs *in vivo* across different eye regions of the zebrafish retina. Surprisingly, this revealed that despite being highly diverse – for example in terms of kinetics and polarity - ACs were mostly non-opponent and spectrally resembled linear combinations of UV- and red-cone signals, which in zebrafish are associated with greyscale processing^7^. Next, we imaged BC light responses in the presence and absence of AC-mediated inhibition. This demonstrated that while BC greyscale processing was profoundly altered across the entire retina, colour processing was affected differently in different regions of the retina. While in the central retina ACs served to set up a new “UV:yellow” axis of colour opponency, in the peripheral retina the population representation of spectral contrast^5,30^ was essentially invariant to the removal of inhibitory signals. However, this was not because opponency in individual BCs was invariant to AC-block. On the contrary: ACs both routinely abolished and generated spectral opponency at the level of individual BCs, but they did so in approximately equal measure, such that the net change across the population of BCs was essentially zero. To preserve the balance between different chromatic and achromatic channels, ACs act near-exclusively through On-circuits.

We conclude that distinct circuits serve to conserve the parsing of colour information performed in the outer retina as the visual signal is transmitted to central and peripheral BCs. The inhibitory interactions within the inner retina that underly other visual processing tasks do not notably alter the population representation of colour information.

## RESULTS

### Surveying amacrine cell functions

To investigate how inhibitory microcircuits in the inner retina contribute to processing we began by recording how the population of amacrine cells (ACs) in larval zebrafish encode greyscale and colour information across the inner plexiform layer (IPL) and across different parts of the eye. For this, we used *in vivo* 2-photon imaging of SyGCaMP3.5 expressed under the ptf1a promoter which targets the vast majority of ACs in zebrafish^25,31^ (Figure 1A-F). We recorded dendritic calcium responses of ACs to a battery of widefield light stimuli testing basic visual processing tasks (Methods): (i) an achromatic (“white”) step of light (3 s On, 3 s Off, 100% contrast) testing response polarity and kinetics (Figure 1D); ii) a frequency modulated chirp centred at 50% contrast testing frequency response (Figure 1D), (iii) steps of light (2 s On, 2 s Off, 100% contrast) at four different wavelengths (‘red’: 592 nm; ‘green’: 487 nm; ‘blue’: 420 nm; ‘UV’ 382 nm) testing spectral sensitivity (Figure 1E) and (iv) ‘tetrachromatic binary noise’ (5 mins, 6.4 Hz, 100% contrast) which allowed us to extract four ‘spectral sensitivity kernels’ per terminal to probe for spectral opponency (Figure 1F, Methods).

**Figure 1.**
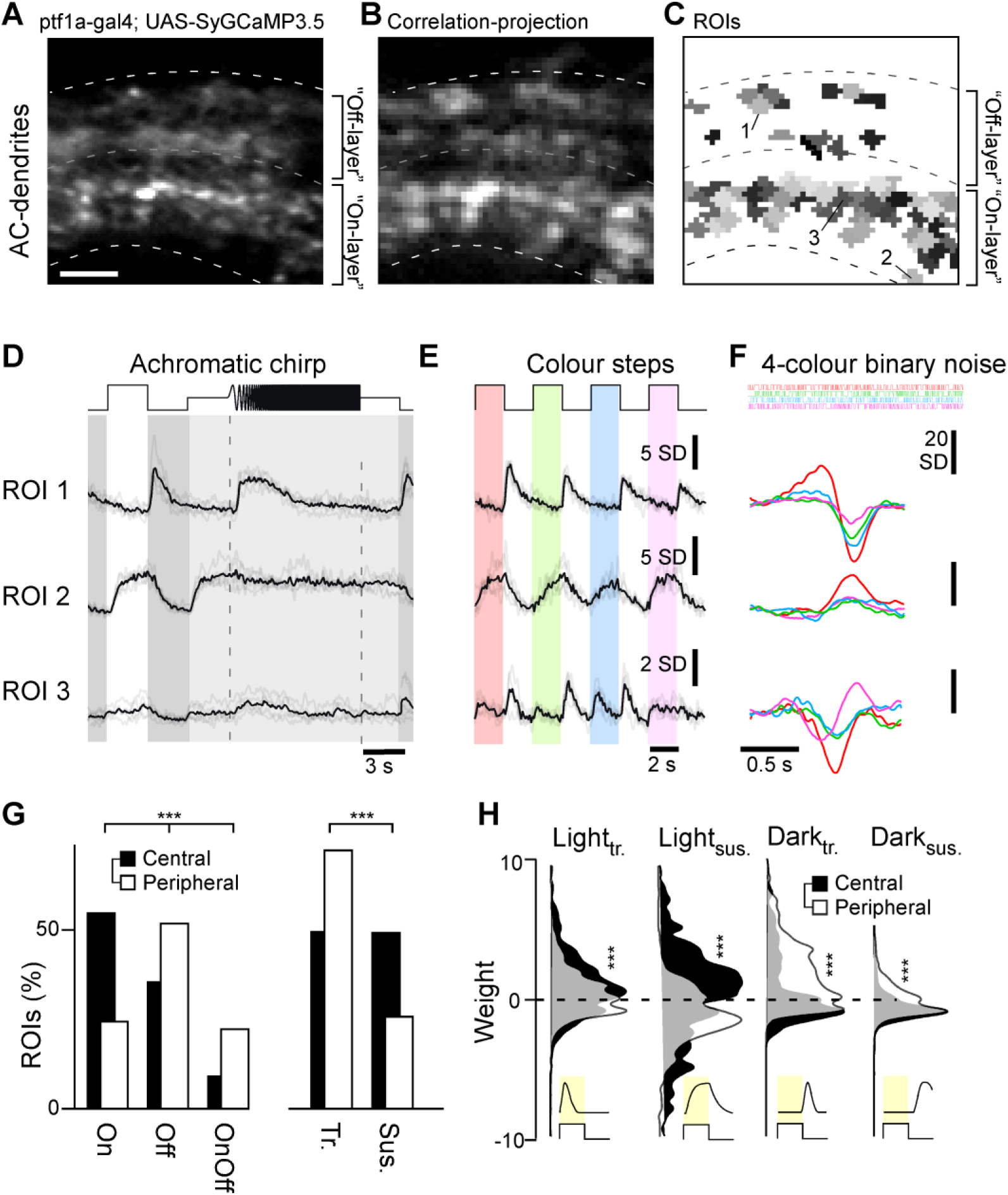
Recording AC-functions in vivo. **A-C**, Example scan of syGCaMP3.5 expressing AC-dendrites within the IPL, showing the scan average (A), a projection of local response correlation, indicating regions of high activity (B) and the correspondingly placed ROI-map (C). **D-F**, Example ROIs (cf. C), with mean trial-averages (black) superimposed on individual repeats (grey) to the three light stimuli tested: an achromatic chirp stimulus (D), four spectrally distinct steps of light from dark (red, green, blue, UV, as indicated, Methods) (E) and a spectral noise stimulus used to establish linear filters (kernels) at the same four wavelengths (F). **G**, Distribution of ROIs from the central (black, n = 927 ROIs, 10 scans, 6 fish) and peripheral retina (white, n = 816 ROIs, 10 scans, 6 fish) by polarity (On/Off/On-Off) and kinetics (transient/sustained). Chi-squared test, p < 0.001 for both datasets. **H**, Distribution of kinetic component weights (Methods, cf. Supplemental Figure S1B) of central and peripheral ROIs. Wilcoxon Rank Sum test, p < 0.001 for all four datasets.

This set of stimuli was chosen to facilitate comparison with previous work^7,13,19,32,33^, and to test a wide range of achromatic and spectral processing tasks within a limited recording time. We recorded from two regions of the eye: (i) the central ‘fovea like’ *area temporalis* which surveys upper frontal visual space to support binocular vision and including prey capture^19,34–38^, and (ii) the peripheral nasal retina, which surveys the outward horizon (n = 10 scans each). ROIs were detected at all IPL depths, with most AC-responses occurring in two major bands, towards the respective centres of the traditional On- and Off-layers (Supplemental Figure S1A). In total, we recorded from n=927 ROIs in the central retina and n=816 ROIs from the peripheral retina.

AC processes exhibited diverse responses across the tested battery of stimuli (Figure 1D-F). ROI 1, for example, consistently exhibited transient Off-responses to both the white chirp stimulus (Figure 1D) and when probed with colour flashes (Figure 1E), while ROI 2 exhibited sustained On-responses. These different behaviours were also captured by the spectral kernels (Figure 1F), which additionally highlighted an overall preference for long-over short-wavelength stimulation in both cases. In contrast, ROI 3 responded poorly to the ‘white’ chirp but exhibited a variety of both On- and Off-responses to different wavelength colour flashes. In this case, the spectral kernels indicated a gradual shift from long-wavelength Off-dominance to short wavelength On-dominance, identifying this ROI as colour-opponent.

### Variations in amacrine cell function in the central and peripheral retina

The larval zebrafish retina is structurally and functionally asymmetrical to acknowledge statistical symmetries in the natural visual world as well as species specific visual demands^19,34,35,39,40^. For example, the central retina comprises a high density of UV-On circuits to support prey capture^19,33,34,41^, while the nasal and dorsal parts of the eye disproportionately invest in colour-opponent circuits^18,19^ to survey the colour-rich horizon and lower visual field, respectively. To evaluate how ACs might contribute to regional differences in the distribution of functional microcircuits, we began by analysing responses to the white step of light, fitting these to a linear kinetic model as described in recent work^18^. Briefly, the model used four kinetic templates to capture the dominant response waveforms across our datasets: Light-transient, Light-sustained, Dark-transient, and Dark-sustained (Supplemental Figure S1B, Methods). This procedure simplified complex responses into four components and their corresponding weights. For example, ROI 2 (Figure 1D) exhibited a relatively slow On-response that was readily captured by a positively weighted Light-sustained component alone (Supplemental Figure S1B, bottom). The Off-response of ROI 1 was kinetically more complex. Capturing this compound waveform required the combined use of a small negatively weighted Light-sustained component alongside a large positively weighted Dark-transient component (Supplemental Figure S1B, top). The same approach served to fit all ACs achromatic step-responses, consistently capturing >93% of the variance across response means.

The extracted component weights allowed us to quantitatively compare the distribution and types of responses across the central and peripheral retina based on two fundamental properties – polarity (On, Off, On-Off) and kinetics (transient, sustained, Figure 1G, Methods). The central retina was On-dominated (54.5% of ROIs, compared to 35.6% Off and 9.4% On-Off ROIs), while the peripheral retina was Off-dominated (51.7% of ROIs, compared to 24.5% and 22.4% of On and On-Off ROIs, respectively; 1.4% non-responders). Moreover, while the central retina comprised approximately equal fractions of transient (49.8%) and sustained (49.5%) responses, the peripheral retina was heavily dominated by transient (72.6%) responses. Further insights emerged by comparing the distributions of kinetic component weights across retinal regions (Figure 1H). Central ACs were dominated by large positive and negative Light-sustained weights, alongside a smaller contribution from positive Light-transient weights. In contrast, peripheral ACs drew on a more varied distribution of kinetic components, and with a particular emphasis on the use of positive Dark-transient weights. ACs from the two retinal regions therefore exhibited a variety of responses to a simple achromatic step of light.

We next sorted ACs into response types based on the full range of stimuli by using a Mixture of Gaussians model to jointly cluster all ROIs, irrespective of IPL position or eye region (Methods). This returned 27 clusters which further highlighted the striking functional differences in AC functions across the eye (Example clusters shown in Figure 2A-E, complete overview in Supplemental Figure S2 and Supplemental Figure S2 Extended 1). Regional information was not used to drive the clustering (see Supplemental Figure S2 Extended 2 for an alternative clustering by eye region) but 20 of the 27 clusters nonetheless comprised ROIs exclusively from either the central or the peripheral retina. The remaining seven clusters comprised various mixtures of ROIs from both regions.

**Figure 2.**
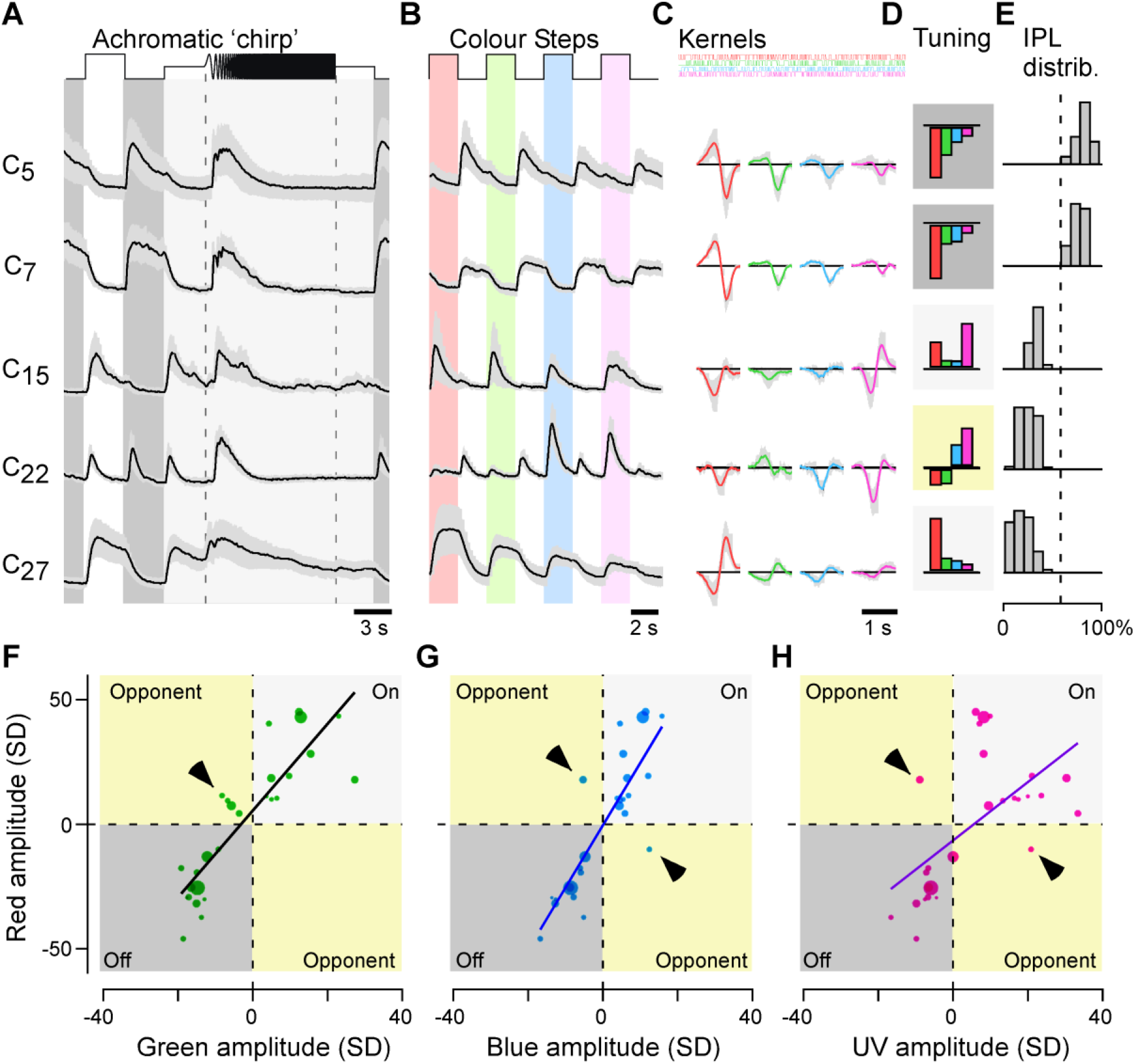
Amacrine cells are kinetically diverse but spectrally simple. **A-E**, Example cluster means±SD in response the white chirp (A) and colour-step stimuli (B), spectral kernels (C) corresponding mean kernel amplitudes (D), and each cluster’s distribution across the IPL (E). Shadings in (D) annotate cluster polarity (dark: Off; light: On, yellow: colour-opponent). **D-H**, Relationship of each clusters’ kernel amplitudes in red (y-axis) plotted against the corresponding amplitudes in green (F), blue (G) and UV (H). Note that most points scatter across the two non-opponent quadrants of the plots, with exceptions highlighted by arrowheads. Dot size indicate the number of ROIs in a cluster. Non-weighted line-fits are superimposed for illustration.

### Amacrine cells are kinetically diverse but spectrally simple

Using the output of the joint clustering procedure, we compared how ACs encode greyscale and colour information. As we detail below, this revealed that ACs could be divided into two main spectral groups: a majority of kinetically diverse but spectrally simple achromatic ACs, and a minority of colour-opponent ACs with complex spectral responses, which stratified exclusively in the On-layer.

21 of the 27 clusters responded similarly to the white and coloured steps of light (Figure 2A,B), a behaviour that was also captured in the spectral kernels (Figure 2C,D). For example, cluster C_5_ exhibited transient Off-responses to steps of light at any wavelength, while cluster C_7_ consistently displayed sustained Off-responses. Correspondingly, each of the four spectral kernels also indicated Off-behaviour (Figure 2C), rendering these clusters non-opponent. Similarly, kinetically distinct On-clusters C_15_ and C_27_ were also non-opponent. The remaining six clusters were more complex. For example, C_22_ displayed transient On-Off responses at all tested wavelengths, however with a notable Off-dominance during long-wavelength stimulation and On-dominance during short-wavelength stimulation, rendering this cluster colour-opponent overall. Such wavelength-dependent rebalancing of On- versus Off- amplitudes – rather than a ‘classical’ full polarity reversal as observed in cones^7^ and bipolar cells^18,19^ – also rendered the remaining five clusters weakly opponent overall.

The dominance of non-opponent responses amongst AC-clusters was further illustrated by comparison of kernel amplitudes across different wavelengths (Figure 2F-H). For example, pairwise comparison of red-versus green-kernel amplitudes highlighted that most clusters exhibited same-sign behaviour at both wavelengths (Figure 2F). Only a minority were red-green opponent (arrowhead), and these clusters notably also exhibited the lowest kernel amplitudes overall. Qualitatively similar behaviour was observed when comparing red-blue and red-UV wavelengths (Figure 2G,H).

The 27 AC-clusters could be sorted into four spectral groups (Methods): Three large non-opponent groups (Off-long biased: C_1-5,7,8,10,16,20,25_; On-long-biased: C_12,14,19,26,27_; “V-shaped”: C_6,9,11,15,17_) and one opponent (RG/BU: C_21,22_; RBU/G: C_13,18,23,24_) (Figure 3A-D). This simplification illustrated their remarkable spectral homogeneity, and further facilitated summarising their distributions across the IPL (Figure 3E-H): All eleven Off-clusters, as well as five of the ten On-clusters followed a common, long-wavelength biased and non-opponent spectral tuning function (Figure 3A,B). Off-long-biased ROIs were mostly, though not exclusively located in the IPL’s traditional Off-layer (Figure 3E), while On-long-biased ROIs were only found in the On-layer (Figure 3F).

**Figure 3.**
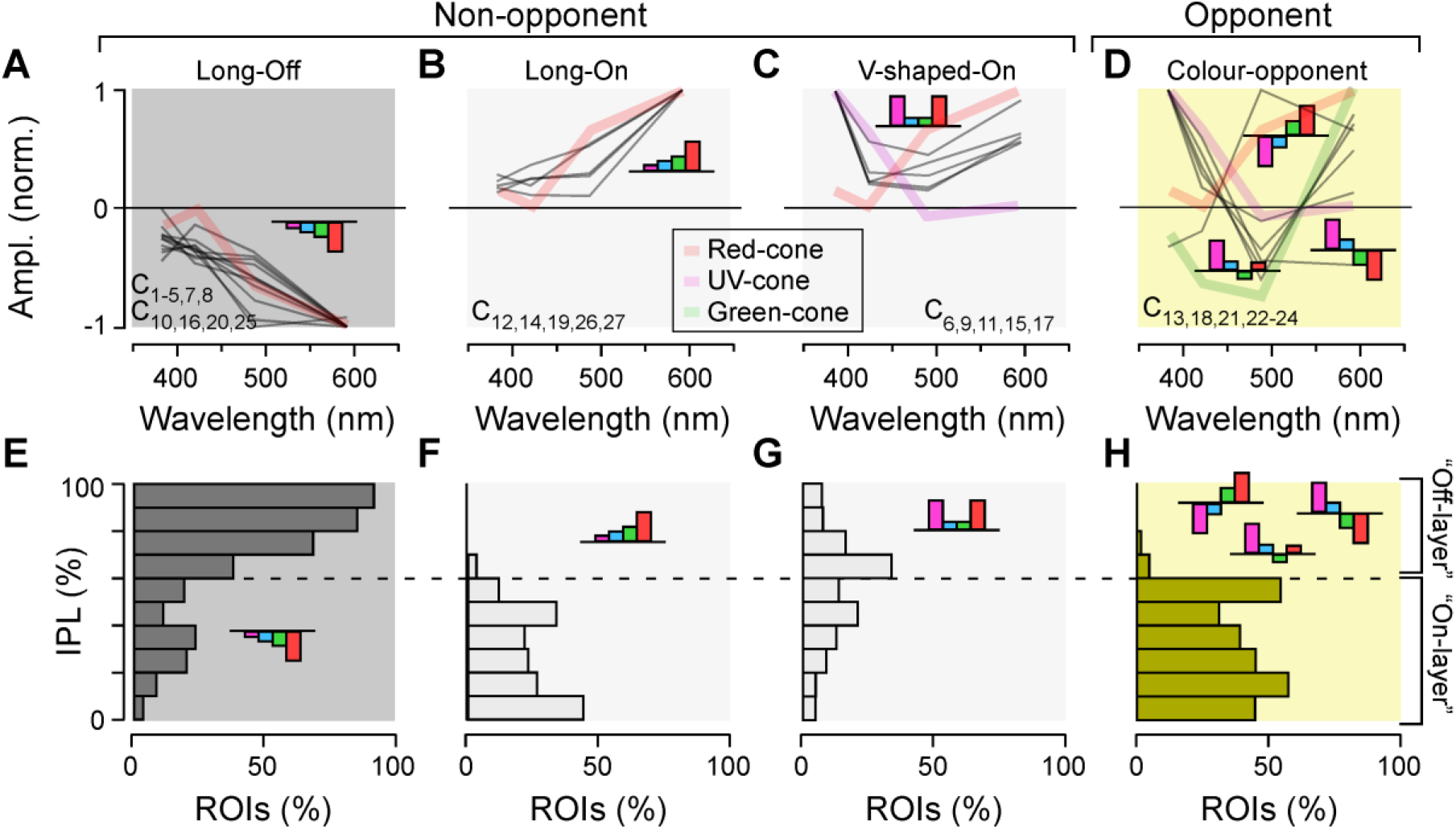
Amacrine cells are kinetically diverse but spectrally simple. **A-D**, Spectral tuning functions of all AC-clusters (grey lines), allocated to one of four groups as shown: Long-wavelength biased Off- (A) and On- (B), V-shaped-On (C), and colour-opponent (D). Plotted behind the AC-spectral tunings are reduced tuning functions of selected cones (cf. Supplemental Figure S3B,C) to illustrate qualitative spectral matches between cones and AC-clusters. Clusters contributing to each group are listed in each panel. **E**,**H**, Corresponding IPL-distribution of AC-ROIs allocated to each of the four spectral groups, as indicated.

All remaining non-opponent On-clusters fell into a third group that was spectrally “V-shaped” (Figure 3C), with ROIs exhibiting an incomplete bias to the centre of the IPL. This last group was overwhelmingly comprised of ROIs from the central retina (Supplemental Figure S3A) which is known to be heavily UV-dominated^18,19,33,34^. The spectral tuning of all 21 non-opponent ACs clusters was readily explained by inputs from red- and UV-cones (Supplemental Figure S3B-E), which are associated with achromatic processing^7^. Strikingly, essentially all ROIs in the Off-layer and approximately half in the On-layer were of this non-opponent population. The remaining half of ROIs in the On-layer consisted of six AC clusters that were both kinetically and spectrally complex, and weakly but consistently colour opponent (Figure 3D,H). The spectral behaviour of these colour-opponent clusters could not generally be explained without additional inputs from the opponent^7^ green-cones which are associated with colour processing (Supplemental Figure S3B-E).

We next tested how these different distributions of amacrine cell functions across the eye and IPL might be linked to spectral and temporal processing in bipolar cells.

### Bipolar cell signalling in the presence and absence of inhibition from amacrine cells

To investigate the effects of AC-mediated inhibition on the visual signal transmitted through the inner retina, we combined pharmacology with *in vivo* 2P imaging of BC synaptic terminals expressing the calcium biosensor SyjGCaMP8m^18,42–44^ (Figure 4A-F, Methods). In each experiment, we first scanned ∼10° eye regions comprising typically 100-120 individual BC terminals (Figure 4A,B) and presented the same battery of stimuli previously used to characterise amacrine cells (Figure 4C-E). Next, we injected a cocktail of gabazine, TPMPA, and strychnine into the eye to pharmacologically block GABA_A_, GABA_C_ and glycine receptors, respectively^26^ (Methods), which represent the major known sources of AC-mediated inhibition in the inner retina^15^. We then imaged the inner retina a second time (e.g. Figure 4F-H) to compare the functions of BC terminals in the presence or pharmacological absence of AC-mediated inhibition.

**Figure 4.**
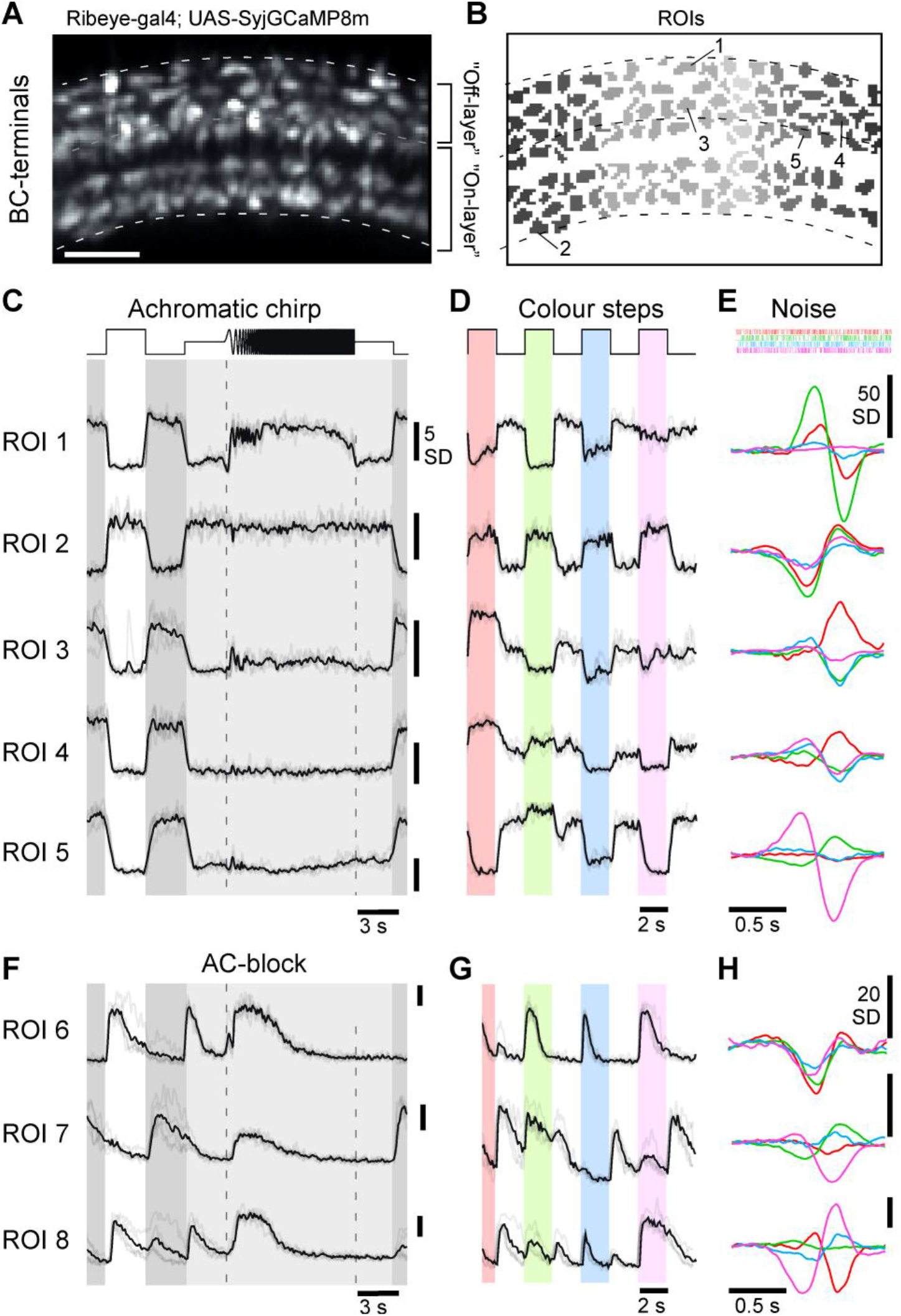
Effects of AC-blockage on BCs. **A**, Example scan field (A) and ROI-mask (B) of a typical scan from BC-terminals expressing SyjGCaMP8m^44,62^, with approximate IPL-boundaries and On/Off layer separation indicated. Scalebar: 5 μm. **C-E**, Example ROIs from (A,B) as indicated, responding to the white chirp (C) and colour-steps (D), alongside mean spectral kernels recovered from 4-colour noise stimulation (E). **F-H**, (As C-E, respectively), but for three different example terminals that were recorded after pharmacological injections of GABAzine, TPMPA and strychnine to block inhibitory inputs from ACs. Note that responses are generally larger and more transient compared to control conditions, but diverse spectral opponencies persist. Scalebars (C,D,F,G): 5 SD, (E): 50 SD, (H): 20 SD.

The efficacy with which this manipulation blocked inhibition in the inner retina was evidenced by the increase in the gain of responses in BC synapses (Figure 5, Supplemental Video 1) and the decorrelation^13^ of these responses (Supplemental Figure S4A,B). We also evaluated the effect of blocking inhibitory receptors on outer retinal function, where horizontal cells spectrally retune the cone output^7^. Using existing cone-type specific SyGCaMP6f lines^7^, we confirmed that the cones’ spectral tunings were invariant to the application of the drug-cocktail (Supplemental Figure S4C-N).

**Figure 5.**
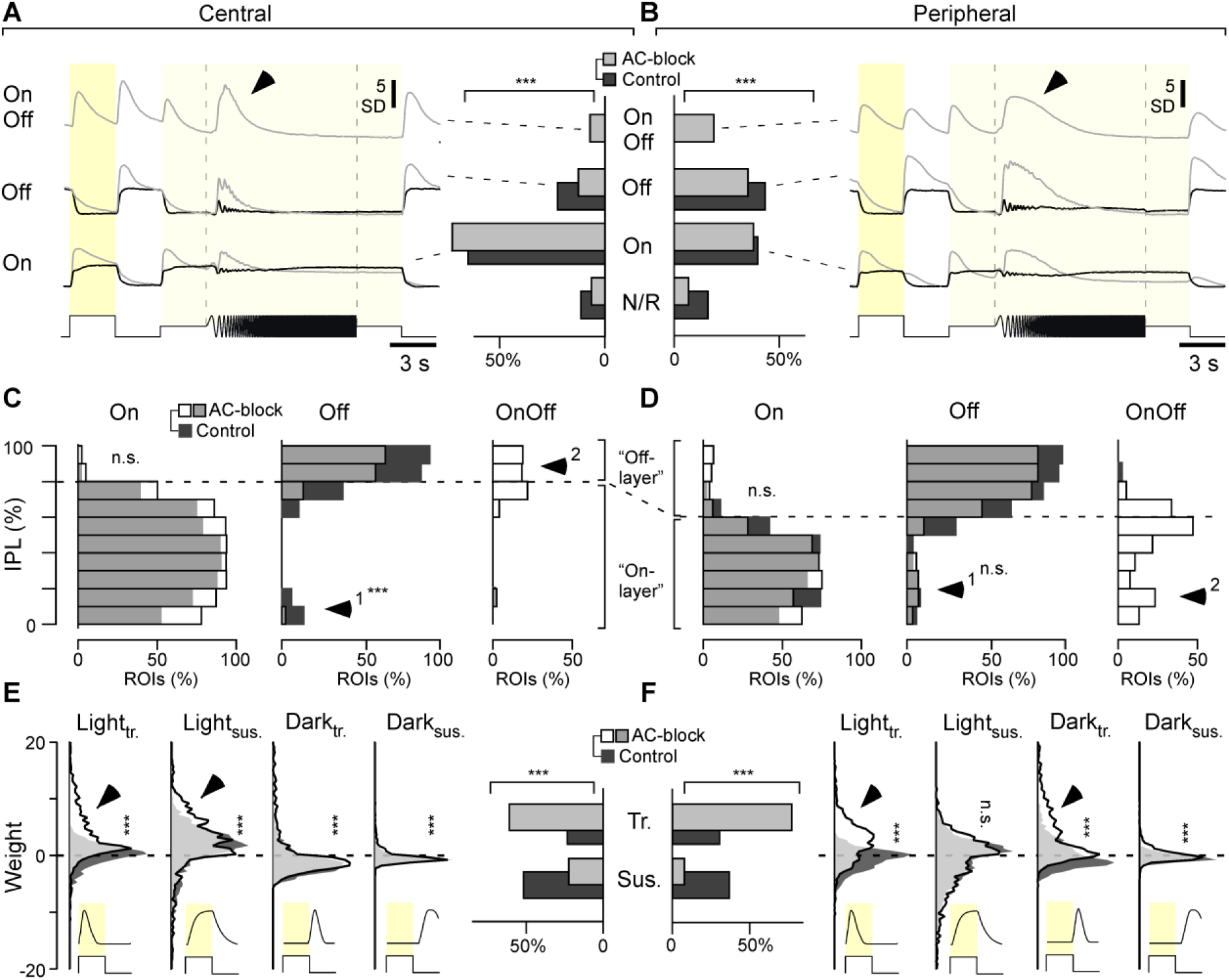
ACs differentially modulate greyscale processing in BCs across the eye. **A**,**B**, Bar plots: Distribution of BC-ROIs recorded in central (A) and peripheral retina (B) by response polarity (Non-responsive, On, Off, OnOff) before (control, dark grey) and after blocking inhibitory inputs (AC-block, light grey) as indicated (Chi-squared tests for the bar plots, p < 0.001 for both datasets), and the corresponding mean response waveform to the white chirp stimulus (cf. Figure 4C) of all terminals in a given category. Arrowheads highlight distinct temporal dynamics during temporal-flicker across the regions. **C, D**, Distribution of On, Off and On-Off BC-terminals across the IPL as indicated with control data shown in dark grey and AC-block data superimposed semi-transparent in white, such that overlapping bars appear in light grey. Note that the On/Off boundary differs between the central (C) and peripheral retina (D, see also Refs^18,19,33^). Arrowheads highlight key differences between central and peripheral histogram-pairs. Two-sample Kolmogorov–Smirnov test, Central: On, p > 0.05; Off, p < 0.01; Peripheral: On, p > 0.05; Off, p > 0.05. Due to the absence of datapoints in control conditions, no statistical tests were used to test changes for OnOff distributions. **E, F**, Distribution of kinetic component weights used to fit BCs’ white step responses (cf. Supplemental Figure S1B, Methods) across the datasets and conditions as indicated (vertical histograms), and distribution into overall transient and sustained response-types (bar plots). Arrowheads highlight the major differences across control and AC-block condition in each case. Wilcoxon rank sum test for the distribution plots, p > 0.05 for peripheral Light_sus_ group, p < 0.001 for all other groups; Chi-squared test for the bar plots, p < 0.001 for both datasets.

In line with previous work^18,19,45^ BCs displayed a broad range of response properties under control conditions, which included both On- and Off cells with diverse temporal and spectral tunings. For example, in a scan from the peripheral retina (Figures 4A,B) ROIs 1 and 2 displayed largely achromatic Off- and On-responses, respectively, while ROIs 3-5 exemplified different forms of colour opponency (Figure 4C-E, Supplemental Video 2). A substantial degree of functional diversity in BC-responses was also observed following pharmacological AC-block, including the continued presence of numerous colour opponent responses (Figure 4F-H). However, the nature and distribution of these disinhibited responses were profoundly altered compared to control conditions, and in a manner that systematically differed between the central and peripheral retina, as we describe below.

### Changes in greyscale processing caused by blocking inhibition

The most general effects of blocking inhibition from ACs were to make responses in BCs larger and more transient, and this occurred across retinal regions and for terminals of all polarities (Figure 5). By fitting step responses to the same four kinetic components previously used to fit ACs we could account for >94% of the variance across BCs. Using the kinetic weights, we automatically classified each BC-response as either unresponsive, On, Off, or On-Off, and computed the average chirp-response traces for the latter three categories per retinal region and condition (Figure 5A,B). In both the central and peripheral retina, blocking ACs reduced the number of Off- and unresponsive terminals and unmasked the presence of ‘intrinsically On-Off’ terminals. Following block of AC inputs, On-Off terminals were also observed in response to coloured stimuli, most notably to red- and UV (Supplemental Figure S5A), but they were never observed under control conditions.

Other effects of blocking inhibition were dependent on retinal region. Unmasking of On-Off responses, for example, was much more common in the peripheral compared to central retina. Moreover, on average, peripheral BCs of all polarities followed the frequency-accelerating part of the chirp for longer compared to central BCs, suggesting regional differences in the modulation of temporal processing. Overall, while blocking ACs mainly accentuated the pre-existing On-bias of the central retina, the same manipulation yielded a more complex re-distribution of response properties in the periphery.

To analyse how these changes in BC function were distributed across the IPL, we segregated terminals into ten strata and computed histograms summarising the relative depth-distributions of On-, Off, and On-Off terminals in each region and condition (Figure 5C,D). Blocking inhibition from ACs had distinct effects in the central and peripheral retina. In the central retina, ectopic Off-responses in the On-layer were abolished (Figure 5C, arrowhead) but these were not affected in the peripheral retina (Figure 5D, arrowhead 1). In the central retina, blocking inhibition also generated mixed On-Off responses in the Off-layer (Figure 5C, arrowhead 2), while in the peripheral retina On-Off response appeared throughout the IPL (Figure 5D, arrowhead 2). These results demonstrate that ACs do not simply regulate the gain and kinetics of the output from BCs, but also the polarity. In the absence of inhibition, 19.3% of BC terminals in the peripheral retina signal both On and Off transitions and these were predominantly in the On-layer. This previously unrecognized function of ACs - regulating the polarity of synaptic activity in BCs - was less prominent in the central retina, where it was only evident in 7.6% of terminals, and these were predominantly in the Off-layer.

Regional differences in the way that ACs interact with BCs were also evident in the temporal domain (Figure 5E,F). Responses tended to become more transient after blocking inhibition but this effect was much stronger in the periphery, where it involved an accentuation of both Light- and Dark-transient response components (Figure 5F, arrowheads). In contrast, kinetic changes in the central retina were more moderate and restricted to Light-transient and Light-sustained components (Figure 5E, arrowheads).

These results demonstrate that ACs interact with BCs in a highly regional manner. In the central retina, ACs regulate the gain and speed of responses, suggesting that here, ACs primarily serve as a gain-control system^46^. But in the peripheral retina, ACs also regulate the segregation of On and Off signals in the On-layer. Below we ask how these functional reorganizations of the inner retina impact spectral processing.

### Changes in colour processing caused by blocking inhibition

To distinguish changes in wavelength from changes in intensity, circuits for colour vision contrast signals of different photoreceptor systems^5,30^. The resultant colour opponent neurons can be considered the fundamental ‘currency’ of colour vision. In zebrafish, BCs represent three types of spectral opponency (Figure 6A): Long- (“red-green”), mid- (“orange-blue”) and short- (“yellow-UV”), with spectral zero crossings at ∼523, ∼483 and ∼450 nm, respectively^18,19^. Of these, long- (“red:green”) and mid-wavelength opponency (“orange:blue”) is already encoded at the level of green- and blue-cones, respectively^7^, implying that ACs are not necessary to establish these channels in BCs. In contrast, short-wavelength opponency (“UV:yellow”) is only weakly represented in UV-cones^7^, but dominant amongst BCs^18^. The expectation, therefore, is that short-wavelength opponency requires the activity of ACs. This expectation was confirmed in the case of BCs in the central retina but not the periphery.

**Figure 6.**
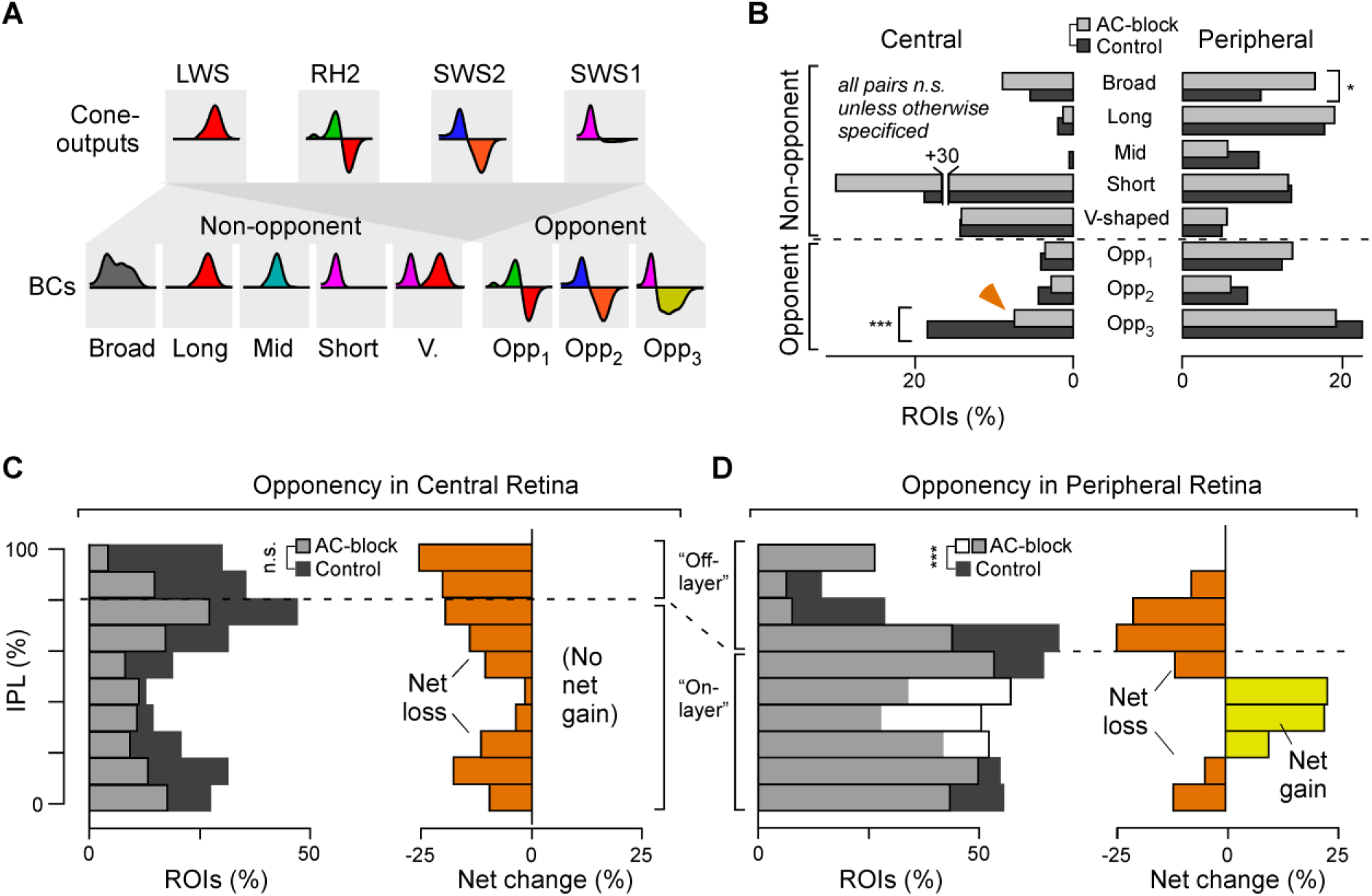
ACs differentially modulate colour processing in BCs across the eye. **A**, Illustration of how cone input could build the different spectral response types in BCs. Insets indicate approximate spectral tuning functions (amplitude y versus wavelength x). The upper row shows the four cones^7^, bottom rows shows BCs^18,19^, divided into five non-opponent and three opponent categories. **B**, Percentages of ROIs per spectral category (cf. A) based on BC kernels (cf. Figure 4E) for central (left) and peripheral datasets (right). Note that the distribution across spectral categories differs strongly by retinal region, but within a region most spectral categories are approximately stable between control (dark grey) and AC-block conditions (light grey). The main exception to this observation is indicated by the arrow. Wilcoxon rank sum tests, central: p > 0.05 for the first 7 groups, p < 0.001 for “Opp_3_” group; peripheral: p = 0.015 for “Broad” group, p > 0.05 for all others. **C, D**, Left: IPL distributions (C, central retina; D peripheral retina) of all ROIs classed as colour opponent during control condition (dark grey) and following AC-block (white, such that their overlay is light grey). Right: Difference between control and AC-block in each case, with net loss of opponency shown in orange, and net gain in yellow. The full datasets leading to the summaries in B-D are shown in Supplemental Data S6. Two-sample Kolmogorov–Smirnov test for changes in IPL distributions of opponent terminals, central: p > 0.05; peripheral: p < 0.001.

Opponency in BCs’ was assessed from spectral kernels computed from colour-noise responses, as used for classifying ACs (Figure 3A-D). However, because BCs^18,19^ are more spectrally diverse than ACs (Figures 1-3), we classified each into one of eight (rather than four) groups: five non-opponent groups (broad, long-biased, mid-biased, short-biased, V-shaped) and three opponent groups (long-opponent: Opp_1_, mid-opponent: Opp_2_, short-opponent: Opp_3_, Figure 6A,B – for full classification see Supplemental Figure S6). This group allocation confirmed previous results^18,19^ that under control conditions, the distribution of spectral response-types strongly differed across the two regions: While in the central retina, spectral groups that required a strong UV-input accounted for the vast majority of responses, the diversity of spectral response types was much more evenly distributed in the peripheral retina, including greater numbers of colour-opponent neurons (peripheral: 43.3%; central: 27.9%). Strikingly, in both retinal regions, these spectral distributions remained largely unchanged following AC-block: No major spectral response group disappeared altogether (Figure 6B, and Supplemental Figure S6), and the abundance of many spectral groups was unchanged. For example, the numbers of V-shaped non-opponent responses appeared entirely unaffected in both retinal regions, while most colour-opponent groups exhibited only marginal changes.

The only significant effect of blocking inhibition on colour opponency that we could detect was a loss of short-opponent responses in the central retina in favour of a corresponding gain in short-biased non-opponent responses (Figure 6B, arrow). This change was not observed in the peripheral retina, where all three opponent groups persisted throughout the pharmacological manipulation.

Having established that the short-opponent interactions between ACs’ and BCs were specific for eye-region, we looked more closely at inhibitory circuits at different locations in the IPL. For simplicity, the distribution of colour opponency was assessed by summing the three colour-opponent groups into a single distribution per experimental condition (Figure 6C,D, individual distributions shown in Supplemental Figure S6B,D). The colour-opponent responses appearing after block of inhibition were both short- and long-wavelength opponent (Supplemental Figure S6D, arrowheads).

This finer analysis revealed that, despite the *overall* numerical conservation of all three opponencies in the peripheral retina following block of inhibition (Figure 6B), this was made up of a loss of opponency in the Off layer and a gain in the On layer (Figure 6D).

### ACs both create and mask colour opponency in individual BCs

To investigate the how blocking inhibition caused a redistribution of colour-opponency in BCs we tracked the *same* terminals across recordings. This approach was not possible when expression of SyGCaMP was driven in all BCs because the high density of terminals in the IPL made it difficult to reliably identify the same terminal before and after the pharmacological manipulation. We therefore performed a new set of experiments using a different transgenic line where BCs expressed SyGCaMP3.5 sparsely (Supplemental Figure S7A). This strategy allowed us to record from 20-30 individual terminals at a time, out of which 40-60% could be reliably matched across control conditions and after blocking inhibition (Supplemental Figure S7A-D). We sampled 182 terminals from 14 fish covering the entire depth of the IPL. Because labelling was sparse, we combined all paired data into a single eye-wide dataset. We also recorded an equivalent but independent sham control dataset (n = 6 scans, n = 144 paired terminals), where we replaced the drug cocktail used to block inhibition with an equivalent volume of non-pharmacologically active vehicle. Sham injections had no significant effect on the functions we analysed (Supplemental Figure S7 Extended, Methods).

As expected, blocking inhibition generally disinhibited BCs, resulting in less selective, more transient, and larger amplitude responses (Figure 7A-E). As in our population dataset, changes in spectral processing were diverse. In ROI-pair 1, responses to colour steps (Figure 7B) were red-biased during control conditions but responded to all four wavelengths after blocking inhibition, and this spectral broadening was also observed at the level of the kernels (Figure 7C). It appears that in this case ACs were masking an intrinsic short-wavelength response to set-up a long-wavelength biased BC. However, the effects on ROI-pairs 2 and 3 were functionally opposite: ROI-pair 2 exhibited a green-UV colour-opponent response during control conditions, which was abolished following AC-block, while vice versa ROI-pair 3 exhibited weak non-opponent response during control conditions but green-UV opponency upon AC-block. Accordingly, in ROI-pair 2, ACs were responsible for setting up BC-opponency, while in ROI-pair 3 ACs masked an intrinsic form of BC-opponency.

**Figure 7.**
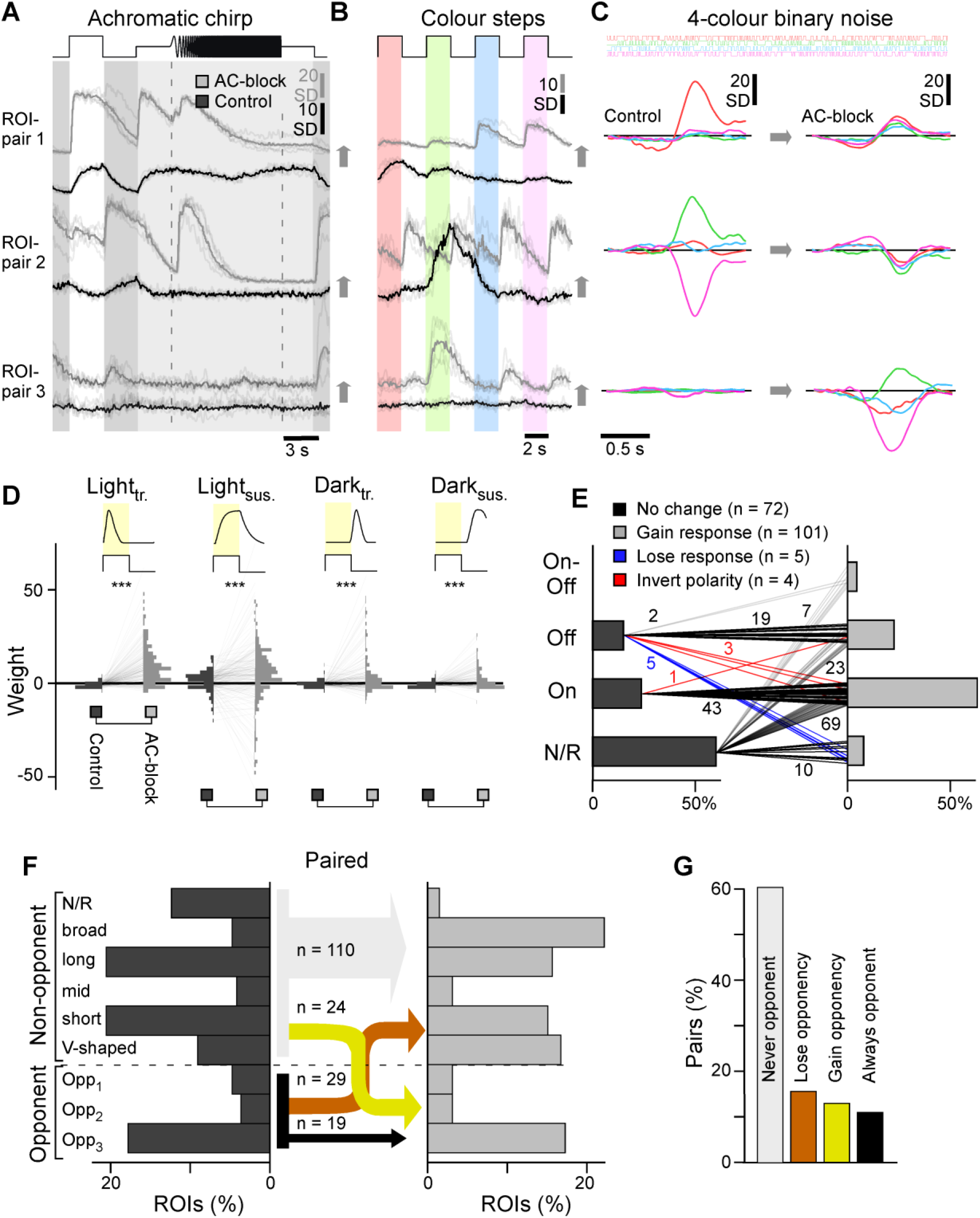
Paired recordings. **A-C**, Example ROI-pairs as indicated in Supplemental Figure S7A-D illustrating frequently observed effects of AC-block on single terminals when probed with the same set of stimuli used to classify all ACs (Figure 1) and BCs (Figure 4). Chirp and colour-step traces (E,F) are vertically offset between the two conditions for clarity. **D**, As Figure 5G,H, but here for paired data (n = 182), pairwise weight distributions for the four kinetic components in control (left) and AC-block condition (right), as indicated. **E**, Paired BCs automatically sorted by polarity in control (left) and AC-block conditions (right). Lines represent the pairs, with numbers of BCs indicated. Chi-squared test, p < 0.001. **F**, As Figure 6B, allocation of BCs into spectral groups, here shown for paired dataset which allowed connecting spectral groups in control condition (left) and following AC-block (right). For simplicity, connections are collapsed into those where there is no change in colour-opponency (grey, black), where colour opponency is lost upon AC-block (orange), and where opponency is gained (yellow), with numbers of BC pairs corresponding to each connection indicated. Note that non-responsive terminals are added as one extra non-opponent category, which resulted from the generally lower signal-to-noise ratio in SyGCaMP3.5 recordings, compared to previously used SyjGCaMP8m. **G**, Summary of changes to colour opponency as shown in (F).

To systematically assess how BC colour opponency is generated and/or destroyed by AC-circuits, we again allocated BC terminals into spectral groups (Figure 7F,G, cf. Figure 6B). Again, there was very little overall change among colour-opponent groups, yet more than half of individual terminals that exhibited colour opponency under control conditions lost their opponency following AC-block (n = 29 of 49, 59.2%). At the same time, an almost equal number of previously non-opponent BCs replenished the population of colour opponent BCs (n = 24). This switching of opponent BCs between conditions affected all three opponent groups: Only 2/8 (25%), 2/7 (29%) and 15/33 (45%) long-, mid- and short-wavelength opponent terminals, respectively, maintained their opponency throughout the pharmacological manipulation. Except for a single BC that switched from long- to short-wavelength opponency, all remaining opponent BCs lost their opponency altogether following AC-block. In fact, for all three colour opponent groups, individual examples could be identified where colour opponency was either preserved (Supplemental Figure S7E-G), lost (Supplemental Figure S7H-J) or gained (Supplemental Figure S7K-L) following AC-block. The overall picture, therefore, is that ACs exert different actions on colour-opponency in different BC terminals: in some terminals, ACs contribute to the generation of colour-opponency but in others they masked pre-existing opponency. These opposing effects of ACs were exerted on all three colour-opponent channels in approximately equal measure.

### ACs modulate BC-spectral processing via the On-channel

The dominance of colour-opponent AC-circuits in the On-layer (Figure 3H) suggests that these ACs have a role in determining how the output from BCs is spectrally tuned. To test this idea, we analysed the colour-step responses of the paired dataset, which included many examples of spectrally selective modulation of On-signals, but spectrally non-selective modulation of Off-signals (Figure 8A-D). For example, under both control conditions and following block of inhibition, the Off-terminal shown in Figure 8A exhibited spectrally broad Off-responses, and ACs acted as achromatic controllers of gain and kinetics. In contrast, the On-terminals shown in Figures 8B-D all exhibited notable spectral changes after block of inhibition. This apparent On-dominance in AC-dependent spectral re-tuning was particularly striking in the example shown in Figure 8D: this BC-terminal exhibited spectrally non-selective Off-responses under control conditions, but block of inhibition unmasked spectrally selective short-wavelength On-responses (arrowheads), making this terminal colour-opponent overall.

**Figure 8.**
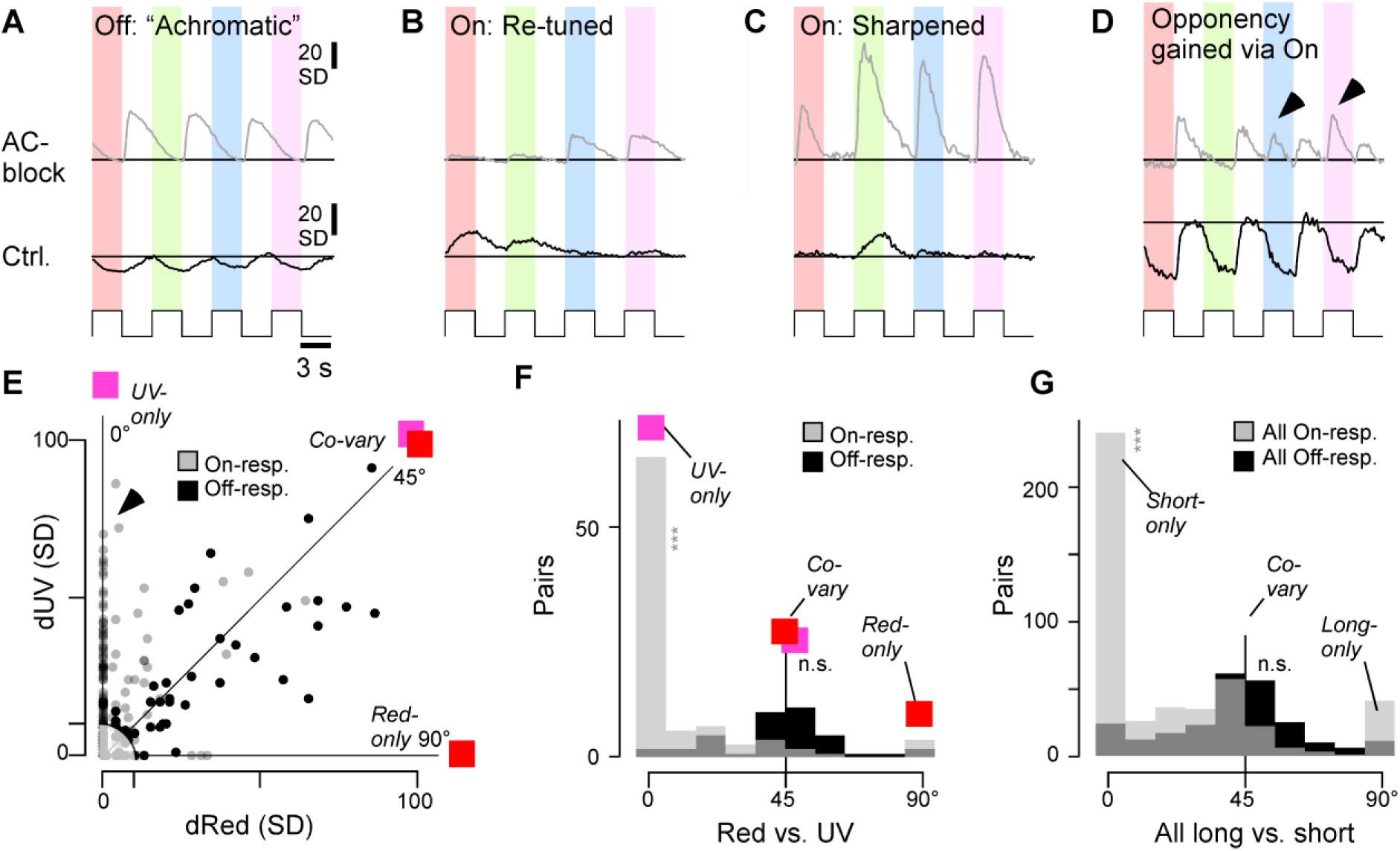
Selective spectral modulation of the On-pathway. **A-D**, Selected example BC colour step responses (trial averages) during control condition (bottom) and following AC-block (top, paired data), illustrating a range of ‘typical’ results. Generally, changes in Off-responses tended to be similar across all tested wavelengths (e.g.: A). In contrast, changes in On-responses tended to be more spectrally diverse, and often effected a change in overall spectral tuning (e.g. panels B,C) including in spectral opponency (e.g. panel D). Arrowheads in (D) highlight wavelength-specific switches in the On-channel that led to a change in colour-opponency. **E-G**, co-variation of absolute response amplitudes changes (in SD) after AC-blockage, compared across different pairs of wavelengths (paired data). (E) shows individual scatterplot for red versus UV. On- and Off-responses plotted separately, as indicated. Correspondingly, an angular histogram (F) was computed from this scatterplot, where 45° indicates co-variation, while peaks around 0° and 90° indicate that one of the two compared wavelength responses changes independently of the other. Datapoints with an Euclidean distance <10 from the origin were excluded from further analysis (shaded area in E). (F) shows the individual red-UV comparison, while (G) shows the sum of all six possible colour combinations (cf. Supplemental Figure S8). Note Off-responses tended to co-vary (peak at 45°), while On-responses exhibited a more diverse distribution, which included peaks are 0°, 45° and 90°. Wilcoxon Signed-Rank test for a given colour combination, tests were performed between On- or Off-angular distributions and 45°: All On (G): p < 0.001, All Off (G): p > 0.05; Red On (F): p < 0.001 (red vs. UV), Off: p = 0.24 (red vs. UV).

To systematically assess the role of On- and Off circuits for shaping chromatic and achromatic circuit functions, responses to each colour-step were again fitted with a weighted sum of four kinetic building-blocks. We reasoned that any achromatic effects of ACs on BCs should lead to a high degree of covariation across wavelengths, while any spectral ‘retuning’ of BCs should manifest in some wavelength-responses being affected more than others. To test this, the degree of response covariation across wavelengths was assessed for each terminal. The plot in Figure 8E shows how blocking inhibition from ACs changed responses in On- and Off-terminals during red stimulation (dRed) plotted against the corresponding amplitude changes in response to UV stimulation (dUV). The equivalence line at 45° is the case where responses to red- and UV covary perfectly. Most Off-responses (black dots) fell near the equivalence line but On-responses showed a mixture of behaviours with a second population falling on or near the 0° line (arrowhead) representing response amplitudes changing to UV steps but not red. Only few points fell on the 90° line, indicating a notable absence of On-responses that were modulated in red without also being modulated in UV.

To summarise this behaviour, we computed the corresponding angular histogram (Figure 8F), which showed a single peak around 45° for Off-responses indicating mostly co-variation, but two main peaks for On-responses: one at 45°, and another at 0°. This general pattern was stable for all possible colour combinations (Figure 8G, individual colour pairs shown in Supplemental Figure S8A-F). In the On-channel, but not in the Off, shorter wavelength responses were modulated more strongly than long wavelength responses.

This analysis provides further evidence that most spectral modulation of BCs by ACs occurs via the On-channel, with short-wavelength circuits being key targets of this modulation.

## DISCUSSION

Investigations of visual processing have usually dealt with stimulus dimensions of space and time, grey-scale processing, separately to colour. Here we have shown that grey-scale and colour processing interact through inhibitory circuits in the inner retina that vary between different zones (Figures 1-3). Blocking inhibition from ACs increased gain in BCs and made responses more transient, as well as unmasking mixed On-Off that were much more common in the peripheral retina compared to the center (Figures 4,5). Simultaneously, ACs contributed to the generation of colour-opponency in the central retina but in the periphery, there was a mixture of effects, enhancing colour-opponency in some BCs while suppressing pre-existing opponency in others (Figures 3,6,7). ACs counteracting intrinsic colour-opponency of BCs acted with a high degree of specificity through just one of the two fundamental channels for grey-scale processing - the On-pathway (Figure 8). We conclude that the central and peripheral retina of larval zebrafish employ fundamentally distinct inhibitory circuits to control the interaction between greyscale- and colour-processing.

### The role of ACs in colour vision

In larval zebrafish, two forms of colour-opponency are established at the level of cone outputs^7^, but three are observed at the level of the downstream BCs^18^. Accordingly, the expectation is that the ‘third’ form of opponency (‘UV:yellow’) is set up by ACs. This was found to be the case in the central retina (Figure 6B,C) but in the periphery the population representation of colour-opponency was remarkably invariant to pharmacological removal of inner retinal inhibition (Figure 6B). At the level of *individual* BCs, however, ACs could either enhance or suppress colour opponency. These opposing effects were balanced across BCs by a combination of factors. First, a majority of ACs was essentially achromatic (Figure 3A-C), indicating that they do not alter spectral processing in a direct manner. Second, a minority of chromatic ACs appear to implement a switch, by which they mask pre-existing colour opponency in some BCs, while at the same time generating qualitatively equivalent information elsewhere (Figure 6D, 7F,G). This switch was implemented mostly by On-circuits (Figure 8), and correspondingly the dendrites of ACs that exhibit spectral opponency were located in the traditional On-layer (Figure 3H).

While it seems intuitive to suggest that colour opponent ACs underpin colour processing in BCs, it may also be that non-opponent ACs are involved in the same task. In principle, combining a spectrally broad AC with a spectrally narrow BC would lead to opponency, while combining an intrinsically opponent BC with a spectrally narrow or V-shaped AC might abolish the opponency. Indeed, blocking inhibition from ACs affected BCs’ colour opponent signals across the entire IPL, including in the Off-layer where colour opponent ACs are absent.

In the future, the different anatomical distributions of colour coding BCs in the presence and absence of AC-inputs (Figure 6D) may provide an important handle for studying the diverse AC-BC circuits that contribute to this overall spectral balancing. Further, understanding if and how these correlative observations are causally linked will likely require the use of more specific transgenic lines that allow more selectively interfering with specific types of BCs and ACs. The same strategy should also help to decipher those BC circuits where ACs mask a pre-existing opponency.

### A special role of On-circuits in zebrafish colour vision?

Most AC-mediated spectral tuning in BCs – whether leading to changes in opponency or simply a rebalancing of non-opponent spectral tunings – were predominately implemented via the On-rather than the Off-channel (Figures 3H, 6D, 8). This observation adds to a growing body of evidence that zebrafish generally leverage On-rather than Off-circuits to compute diverse aspects of colour-information. For example, both at the level of the retinal output^33^, and within the brain^32,47,48^, most spectral diversity is represented in the On-channel. In contrast, the spectral tuning function of the brain’s overall Off-response essentially resembles the spectral tuning function of red-cones in isolation, which also corresponds with the mean-spectrum of natural light in the zebrafish natural habitat^7,32^. From here, it is tempting to speculate that zebrafish generally use the Off-channel as an ‘achromatic reference’, while On-circuits can, where required, provide spectrally biased points of comparison to serve spectral and colour vision. A predominant use of one rather than both polarities for encoding spectral information could also be advantageous in that it might permit largely unaltered travel of the red-cones’ achromatic signal to the brain: by restricting the bulk of spectral computations to the On-strata of the IPL, circuits within the Off-strata can operate in an essentially achromatic manner. In agreement, the vast majority of ACs in the Off layer exhibited such achromatic tunings (Figure 3A,E). In the future it will be interesting to test if such an On-dominance amongst spectral computations is also a feature in other species.

The possible link between polarity and distinct spectral functions also raises the question how On-Off responses are used for spectral coding. Among ACs, we observed a strong link between colour opponency and On-Off responses (Figure 2B, Supplemental Figure S2). Correspondingly, among BCs, intrinsic On-Off responses were routinely unmasked by blocking ACs, particularly in the central retina where they were present throughout the On-layer. (Figure 5A,B,E,F, Supplemental Figure S5A). In the future it will therefore be important to probe more directly to what extent On-Off processing in ACs and BCs can be causally linked to specific aspects of colour processing. Another unanswered question is the source of BCs’ On-Off responses. Opponency in cones alone is unlikely to explain this observation. This is because cone-inversions from their intrinsic Off-response to an HC-mediated overall On-response occurs exclusively at long-wavelengths^7^; however, many unmasked On-Off BCs were short-wavelength biased (Supplemental Figure S5A). Instead, as for the observed intrinsic UV:yellow opponent BCs (see above), their existence points to the presence of yet unexplained mechanisms of signal transfer between cones and BCs in the zebrafish retina. One putative mechanism might involve On-acting excitatory amino acid transporters (EAATs)^49,50^.

### Colour opponency in the absence of ACs

The complex interplay of masked and generated BC opponencies in the absence of inner retinal inhibition confirms that BCs inherit diverse spectral opponencies from the outer retina^5,7,18^. However, the full picture is decidedly more complex than anticipated from previous work. That long- and mid-wavelength opponencies can be preserved in BCs in the absence of ACs is perhaps expected, since these two axes are already fully represented by the two mid-wavelength cones^7^. However, it remains unclear how the third, short-wavelength opponent axis can persist. While, as with primates^51^, zebrafish UV-cones (SWS1) also exhibit weak but significant “UV:yellow” opponency^7^, it seems implausible that this can account for the observed effects. First, this pre-existing outer retinal opponency would need to be substantially boosted to match the much more pronounced opponency in BCs^18^. Second, in cones, this opponency was restricted to the central retina, and, therefore, it cannot account for the profusion of BCs’ UV:yellow opponencies observed in the periphery^18^.

The continued presence of UV:yellow opponency in BCs following AC-block strongly points to the existence of a mechanism capable of selectively inverting cone signals within single BCs. Three potential and non-mutually exclusive mechanisms present themselves. First, a single BC might express both depolarising and hyperpolarising glutamate receptor systems at their dendritic tips that contact different cones (see also the above discussion on BCs’ On-Off signals). Second, BCs could receive direct inputs from HCs. For example, a putative BC driven by sign-inverted inputs from UV-cones (i.e. UV-On) could simultaneously receive sign-preserving inputs from H1 and/or H2 HCs, which themselves carry a sign-preserving long-wavelength biased signal^7^. In zebrafish, the presence of direct inputs from HCs to BCs has not been observed; however, the concept is tentatively supported by the anatomical presence HC-BC contacts in mice^52^. Third, zebrafish might have ACs that use fast neurotransmitters other than GABA and/or glycine, which presumably continue to function throughout our pharmacological interventions. For example, mice feature VGlut3 ACs, a population of part-glutamatergic ACs implicated in motion processing^53–55^. Similarly, another key neuron implicated in mammalian motion processing is the starburst amacrine cell (SAC) which co-releases acetylcholine alongside GABA^56,57^. However, the functional role of SACs outside mammals remains sparsely explored^58^.

### The role of green- and blue-cone circuits in supporting inner retinal colour processing

Unlike red- and UV-cones, zebrafish green- and blue-cones provide strongly colour opponent outputs due to feedforward signals from the HCs^7^. Accordingly, these cones might directly support colour opponency in BCs. In support of this hypothesis, the spectral zero-crossings marked by these two cones remain represented within BCs, both in the presence and in the absence of ACs. However, only a minority of green- and blue-cone-like BCs retained their specific opponency upon AC-block (Figure 7A). This suggests that while the signals from green- and blue-cones can be directly used to support colour opponency in BCs, this motif is by no means dominant when considering the complete circuit. Instead, most BC circuits that represent these two spectral opponencies required inputs from ACs. In zebrafish, green-but not blue-cones provide cone-type-exclusive drive for at least two anatomically distinct types of BCs^12^, providing a possible neural substrate for the minority of green-cone-like BCs that were unaffected by AC-block. These might account for some of the unmasked green-cone-like BCs when ACs were blocked. Possible green-cone-exclusive BCs might also link with the observation that green-light stimulation could result in long-wavelength biased spectral effects on BCs (Supplemental Figure S9D,E), and that most opponent ACs seemed to be partially built from green-cone inputs (Supplemental Figure S3D,E).

In contrast, the possible roles of blue-cones in zebrafish colour vision remain much more elusive. A blue-cone-exclusive BC is not known to exist^12^, which leaves the origin of any intrinsic blue-cone-like BC-tunings unclear. Further, we found no evidence of any major involvements of blue-cones in AC-processing (Supplemental Figure S3D,E). On the other hand, the four LEDs used in the present study were not optimally placed to disambiguate blue-from UV-cone contributions (Supplemental Figure S3B,C). Nevertheless, our findings add to a perhaps puzzling body of evidence that questions a key role of blue-cones in shaping larval zebrafish vision^5^.

### An evolutionary perspective

These results provide tentative insights into the evolution of computation in the brain: in vertebrates and diverse invertebrate eyes alike, the evolution of colour computations likely preceded the evolution of complex spatiotemporal vision. This is because (i) opsins including their immediate spectral diversification preceded the evolution of highly resolved spatial vision in any animal by some 250 million years^59^, (ii) all extant vertebrates, including lampreys, feature subsets of the same four ancestral opsins across their photoreceptors^4,5^, and (iii) particularly in shallow water where vision first evolved, spectral information provides a wealth of behaviourally critical cues that do not categorically need supplementing with spatial information (discussed e.g. in Refs^5,59,60^). From here, it seems plausible that the earliest forerunners of vertebrate eyes^61^ gradually evolved the bulk of their inner retinal circuits on top of well-functioning outer retinal circuits that already provided useful spectral information. In such a scenario, inner retinal circuit evolution would have occurred under constant selection pressure to maintain coding efficiency for colour vision, thus perhaps explaining the arrangement that we see in zebrafish today. This interpretation would further imply that perhaps also in other layered networks of brains, the primary function of some microcircuits may not be to create new computations, but rather to make up for computations which would otherwise be lost.

## METHODS

### Lead Contact

Further information and requests for resources and reagents should be directed to and will be fulfilled by the Lead Contact, Tom Baden.

### Data and Code Availability

Pre-processed functional 2-photon imaging data and associated summary statistics will be made freely available on Data Dryad and via the relevant links on http://www.badenlab.org/resources and http://www.retinal-functomics.net.

### Materials Availability

The transgenic lines Tg(ribeye: SyjGCamp8m), Tg(ribeye:Gal4; UAS:SyGCamp3.5), Tg(ptf1a:Gal4;UAS:SyGCamp3.5), Tg(opn1sw1:GFP:SyGCaMP6f), Tg(opn1sw2:SyGCaMP6f), Tg(LCRhsp70l:SyGCaMP6f), Tg(trb2:syGCamp6f) used in this study have been previously published^25^ and are also available upon request to the lead author.

## EXPERIMENTAL MODEL AND SUBJECT DETAILS

### Animals

All procedures were performed in accordance with the UK Animals (Scientific Procedures) act 1986 and approved by the animal welfare committee of the University of Sussex. Animals were housed under a standard 14:10 day/night rhythm and fed three times a day. Animals were grown in 0.1 mM 1-phenyl-2-thiourea (Sigma, P7629) from 1 *dpf* to prevent melanogenesis. For all experiments, we used 6-8 days post fertilization (*dpf*) zebrafish (Danio rerio) larvae. For 2-photon *in-vivo* imaging, zebrafish larvae were immobilised in 3% low melting point agarose (Fisher Scientific, BP1360-100), placed on a glass coverslip and submerged in fish water. Eye movements were prevented by injection of α-bungarotoxin (1 nL of 2 mg/ml; Tocris, Cat: 2133) into the ocular muscles behind the eye.

## METHOD DETAILS

### Light Stimulation

With fish mounted on their side with one eye facing upwards towards the objective, light stimulation was delivered as full-field flashes of light. For this, we focused a custom-built stimulator through the objective, fitted with band-pass-filtered light-emitting diodes (LEDs) (‘red’ 588 nm, B5B-434-TY, 13.5 cd, 8°; ‘green’ 477 nm, RLS-5B475-S, 3-4cd, 15°, 20 mA; ‘blue’ 415 nm, VL415-5-15, 10-16 mW, 15°, 20 mA; ‘ultraviolet’ 365 nm, LED365-06Z, 5.5 mW, 4°, 20 mA; Roithner, Germany). LEDs were filtered and combined using FF01-370/36, T450/pxr, ET420/40 m, T400LP, ET480/40x, H560LPXR (AHF/Chroma). The final spectra approximated the peak spectral sensitivity of zebrafish R-, G-, B-, and UV-opsins (Supplemental Figure S3B-D), respectively, while avoiding the microscope’s two detection bands for GFP and mCherry. To prevent interference of the stimulation light with the optical recording, LEDs were synchronized with the scan retrace at 1 kHz (1 ms line duration) using a microcontroller and custom scripts. Further information on the stimulator, including all files and detailed build instructions can be found in Ref^43^.

Stimulator intensity was calibrated to be spectrally flat at 30 mW per LED, which corresponds to low-photopic conditions. Owing to 2-photon excitation of photopigments, an additional constant background illumination of ∼10^4^ R^*^ was present throughout^64,65^. For all experiments, larvae were kept at constant illumination for at least 5 seconds after the laser scanning started before light stimuli were presented. Three types of full-field stimuli were used: (i) a spectrally flat ‘white’ chirp stimulus^66^ where all four LEDs were driven together, (ii) flashes of light at each of the four wavelengths (2 s On, 2 s Off), and (iii) a binary dense and spectrally flat white noise, in which the four LEDs were flickered independently in a known random binary sequence at 6.4 Hz for 300 seconds.

### 2-photon calcium imaging

All 2-photon (2P) imaging was performed on custom-built 2P microscope equipped with a mode-locked Ti:Sapphire laser (Chameleon 2, Coherent) tuned to 915 nm for SyGCaMP imaging. Emitted photons were collected through the objective (Nikon, MRD77225, 25X) as well as through an oil condenser (NA 1.4, Olympus) below the sample using GaAsP photodetectors (H10770PA-40, Hamamatsu). For image acquisition, we used ScanImage software (r 3.8) running under Matlab (R 2013b). All recordings were taken at 128 × 100 pixels (10 Hz frame rate at 1 ms per scan line).

### Pharmacological manipulation

For pharmacological AC blockage, we injected ∼4 nL of a solution containing antagonist to GABA_A_, GABA_C_ and glycine receptors into the anterior chamber of the retina. The estimated final concentration in the extracellular space was 5 μM gabazine (Sigma) as antagonist of GABA_A_ receptors; 5 μM TPMPA (Sigma) as antagonist of GABA_C_ receptors; 5 μM strychnine (Sigma) as antagonist of glycine receptors^26^. The solution also contained 1 mM Alexa 594 for verifying the quality of the injection.

### Data analysis

Data analysis was performed using IGOR Pro 6.3 and 8.2 (Wavemetrics), Fiji (NIH) and Matlab R2019b / R2020b (Mathworks).

### ROI placement, IPL detection and functional data pre-processing

Where necessary, images were xy-registered using the registration function provided in SARFIA^67^ running under IGOR Pro 8.2. For AC data, ROIs were defined automatically based on local image correlation over time, as shown previously^13^. For densely labelled SyjGCaMP8m BC data, regions of interests (ROIs) were placed automatically after local thresholding of the recording stack’s standard deviation (typically >20) projection using the tetrachromatic noise data, followed by filtering for size and shape using custom-written scripts running under IGOR Pro 8.2 (WaveMetrics), as described previously^34^. For sparsely labelled SyGCaMP3.5 BC data, ROIs were drawn by hand based on the standard deviation projection across the tetrachromatic noise data. The ROIs from control condition and drug condition were drawn separately. Terminals were paired across the two conditions using the experimenter’s best judgment, which we found to be more reliable than automated procedures. The matching of terminals across conditions was greatly facilitated by the sparse expression strategy, and throughout we tried to be as conservative as possible to only match terminals when we were certain that they are the same ones (i.e. minimising false positives, at the expense of false negatives). For cone recordings, the ROIs were drawn manually from the standard deviation projection across time.

In all scans within the IPL layer, IPL boundaries were drawn by hand using the custom tracing tools provided in SARFIA^67^. The IPL positions were then determined based on the relative distance of a ROIs’ centre of mass between the IPL boundaries and mapped to the range 0% to 100%.

Fluorescence traces for each ROI were z-normalised, using the time interval 2-6 seconds at the beginning of recordings as baseline. A stimulus time marker embedded in the recording data served to align the Ca^2+^ traces relative to the visual stimulus with a temporal precision of 1 ms. Responses to the chirp and step stimuli were up sampled to 1 kHz and averaged over 5 trials. For data from tetrachromatic noise stimulation we mapped linear receptive fields of each ROI by computing the Ca^2+^ transient-triggered-average. To this end, we resampled the time-derivative of each trace to match the stimulus-alignment rate of 500 Hz and used thresholding above 0.7 standard deviations relative to the baseline noise to the times *t*_*i*_ at which Calcium transients occurred. We then computed the Ca^2+^ transient-triggered average stimulus, weighting each sample by the steepness of the transient:

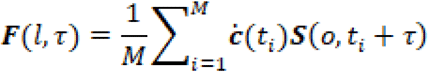

Here, ***S***(*l,t*) is the stimulus (“LED” and “time”), *τ* is the time lag (ranging from approx. −1,000 to 350 ms) and *M* is the number of Ca^2+^ events. The resulting kernels are shown in z-scores for each LED, normalised to the first 50 ms of the time-lag. To select ROIs with a non-random temporal kernel, we used all ROIs that exceeded a standard deviation of ten in at least one of the four spectral kernels. The precise choice of this quality criterion had no major effect on the results.

### Kernel polarity

The use of a fluorescence-response-triggered average stimulus (here: ‘kernel’) as a shorthand for a neuron’s stimulus-response properties, while potentially powerful (e.g. Refs^13,19,33^), ought to be considered with some caution. For example, determining a binary value for a kernel’s polarity (On or Off) can be conflicted with the fact that a neuron might exhibit both On and Off response aspects. Moreover, different possible measures of On or Off dominance in a kernel can generate different classification biases. Here, following our previously established approach^19,33^ we defined On and Off based on a measure of a kernel’s dominant trajectory in time. Before the calculation, we first smoothed the kernels to eliminate the high-frequency noise. After that, we determined the position in time of each kernel’s maximum and minimum. If the maximum preceded the minimum, the kernel was classified as Off, while vice versa if the minimum preceded the maximum, the kernel was defined as On.

### Reconstruction of step responses using kinetic components

To reconstruct each cell’s mean response into constituent spectral and temporal components we used four temporal components associated with a given light response (i.e. 3 s light, 3 s dark for ‘white’ steps, and 2 s light 2 s dark for ‘colour’ steps), following a previously described approach^18^. The temporal components used resembled light-transient, light-sustained, dark-transient, and dark-sustained temporal profiles (Supplemental Figure S1B). These components were fitted to the trial-averaged step responses of individual ROIs in sequence, in each case minimising the mean squared difference between a template’s peak and the corresponding five time-points in measured response, with previously fitted components subtracted. The fit sequence was: Light-sustained, light-transient, dark-sustained, dark-transient. This yielded four corresponding weights, scaled in z-scores in accordance with the amplitudes of the trial averaged response means.

To assess reconstruction quality, reconstructed data was subtracted from the original ROI-means to yield residuals. From here, we compared original data, reconstructions, and residuals based on variance explained across all ROIs (as in Ref^18^). To this end, we first computed the total variance across all clusters for each time-point, computed the area under the curve for each variance-trace, and normalised each to the result from the original cluster means. By this metric, reconstructions captured 98%, 97%, 95% and 96% of the total variance for the ‘white-control’, ‘white-AC-block’ and ‘colour-control’ and ‘colour-AC-block’ steps, respectively.

Response polarity per ROI was then computed as follows. A ROI was considered as displaying an On-response if the sum of the light-transient and light-sustained weights exceeded 3 SD. A ROI was considered as displaying an Off-response if either the sum of the light-transient and light-sustained weights was more negative than −3 SD, or if the sum of the dark-transient and dark-sustained components exceeded 3 SD. If by these criteria a ROI display both On- and Off-responses, it was counted as On-Off. ROIs failing to elicit either On- or Off-responses were counted as non-responsive.

### Clustering of ACs

Clustering was performed on the dataset containing the functional responses of ACs to chirps, colour steps and kernels derived from the colour noise stimulus. All input traces were up sampled to 1 kHz (cf. pre-processing) which yielded n = 25,000 points (chirp), four times n = 4,000 points (steps) and four times n = 1,299 points (kernels). Responses to all three stimuli were used for the clustering.

Regions of interest (ROIs) with low-quality responses to all three stimuli were identified and removed from the data set, ROIs with a high-quality response to at least one stimulus being retained in all cases. The quality of response to the chirp and step stimuli was determined using the signal-to-noise ratio quality index: *QI* = *Var*[⟨***C***⟩_*r*_]_*t*_/⟨*Var*[***C***]_*t*_⟩_*r*_, where *C* is the *T* by *R* response matrix (time samples by stimulus repetitions), and ⟨·⟩_*x*_ and *Var*[·]_*x*_ denote the mean and variance respectively across the indicated dimension, *x* ∈ {*r,t*} (see Ref^66^). A quality threshold of 0.35 was chosen, below which responses were judged to be of poor quality. We calculated the standard deviation in the light intensity over time for each stimulus colour in the kernel (R, G, B and UV). The kernel quality of each ROI was defined as the maximum standard deviation across the four colours. A kernel quality threshold of 5 was chosen, below which kernels were judged to be of poor quality. The raw data set was of size n = 1,776. Following quality control, the data set was of size: n = 1,743 (98.1% of the original).

We scaled the data corresponding to each chirp, step colour and kernel colour by dividing each one by the standard deviation through time and across ROIs. In this way we ensured an even weighting between stimuli.

We used principal component analysis (PCA) to reduce the dimensions of the problem prior to clustering. PCA was performed using the Matlab routine **pca** (default settings). We applied PCA separately to the chirps and to the portions of a data set corresponding to each of the step and kernel colours, retaining the minimum number of principal components necessary to explain ≥99% of the variance. The resulting nine ‘scores’ matrices were then concatenated into a single matrix ready for clustering. The following numbers of principal components were used – chirp: 41; step: 8 R components, 9 G components, 13 B components and 13 UV components; kernels: 7 R components, 16 G components, 31 B components and 21 UV components, giving 159 PCA components in total.

We clustered the combined ‘scores’ matrix using Gaussian Mixture Model (GMM) clustering, performed using the Matlab routine **fitgmdist**. We clustered the data into clusters of sizes 1,2,…,50, using i) shared-diagonal, ii) unshared-diagonal, iii) shared-full and iv) unshared-full covariance matrices, such that (50*4 =) 200 different clustering options were explored in total. For each clustering option 20 replicates were calculated (each with a different set of initial values) and the replicate with the largest loglikelihood chosen. A regularisation value of 10^−5^ was chosen to ensure that the estimated covariance matrices were positive definite, while the maximum number of iterations was set at 10^4^. All other **fitgmdist** settings were set to their default values.

The optimum clustering was judged to be that which minimised the Bayesian information criterion (BIC), which balances the explanatory power of the model (loglikelihood) with model complexity (number of parameters). Lastly, clusters with <10 members were removed.

Using the above procedure, we obtained 23 clusters (1 cluster with <10 members was removed), with unshared diagonal covariance matrices providing the optimal solution. Finally, we split n = 4 clusters with a notably bimodal IPL distribution into their On- and Off-stratifying components (IPL-cut at 60% depth), yielding a total of 27 response groups. To what extent the 27 clusters correspond to AC-types remains unknown, and in view of >60 AC types in mice^22^, 27 putative types in zebrafish probably underestimates their full diversity, which is likely part related to the necessarily incomplete sampling of the full stimulus space and/or possible incomplete labelling of the ptf1a driver.

### Fitting of AC-cluster spectral tuning functions with cones

Spectral tuning functions of AC clusters means were matched to those of previously recorded cones (cf. Supplemental Figure 3B,C) based on the four relative kernel amplitudes (as shown in Figure 3A-D). Fitting was done as follows: For each tested cone-combination (e.g. all cones, or R+U only, etc.) we computed 10^6^ possible combined tunings at random by summing the respective reduced cone tuning functions (Supplemental Figure 3C) with random scaling between −1 and 1 each. We then computed the linear correlation coefficient between each AC-cluster’s tuning function, and each of the randomly generated combined cone-tunings, in each case choosing cone-combination that gave the maximal correlation as the best fit. Finally, for each best fit, we scaled all cone weights to minimise the residual between the corresponding AC-cluster’s tuning function and that of the combined cone-tuning. Based on this strategy, the mean variance explained for cone-combinations RGBU, RU, RGU, GU, respectively, were: All long-On/Off clusters: 95.4%, 86.3%, 90.0%, 53.3%; All V-shaped clusters: 84.0%, 81.4%, 84.9%, 55.6%; All opponent clusters: 86.8%, 63.2%, 83.1%, 71.8%.

### Sorting ACs and BCs into spectral groups

Using the amplitudes of the four kernels, cluster means of ACs were sorted into one of the four spectral groups. First, the AC clusters were divided into non-opponent (Figure 3A-C) and opponent groups (Figure 3D). Next, For the non-opponent groups, the AC clusters were sorted into the three following groups: Long-Off, with the long-biased Off-response across all four spectral stimuli if *abs*(R+G)>*abs*(B+U)*2; Long-On, as Long-Off but for On-responses; V-shaped: *abs*(R+U)>*abs*(G+B)*2. The opponent group was further sorted into RG/BU (C_21,22_) and RBU/G (C_13,18,23,24_) sub-groups based on the relative signs of green vs. UV-responses.

Kernel amplitudes of each BC terminals were sorted into 8 spectral groups. First, all BCs that failed to exceed a minimum absolute amplitude of 10 SD in at least one of the four kernels was counted as non-responsive. Next, we divided the remaining BCs into non-opponent (groups 1-5) and opponent types (groups 6-8) based on the relative signs of all four (R, G, B, U) kernels. From here, the two sets of BCs were sorted further as follows: *Non-opponent BCs*, in order: Long-biased if *abs*(R+G)>*abs*(B+U)*2, Short-biased if *abs*(R+G)<*abs*(B+U), Mid-biased if *abs*(G+B)>*abs*(R+U)*2, V-shaped if *abs*(R+U)>*abs*(G+B)*2, else: “Broad”. Opponent BCs; Short-opponent if (R+G>0 && U<0) || (R+G<0 && U>0), Mid-opponent if (R>0 && B<0) || R<0 && B>0), Long-opponent if (R>0 && G<0) || (R<0 && G>0). Finally, in rare cases where (R>0 && G<0 && U>0) || (R<0 && G>0 && U<0), BCs were allocated as long-opponent if *abs*(R)>*abs*(U) but as short-opponent *abs*(R)<*abs*(U).

Note that despite the similar sorting procedure, ACs only fell into 4 groups (Figure 3A-D) while BCs can be sorted into 8 (Figure 6B).

### Response Synchronisation

To determine the degree of response synchronisation within each scan, we used the synchrony measurement method described in Ref^63^, using the z-normalized fluorescence traces from tetrachromatic noise stimulation of the wide-expressed syGCaMP8m BC data as the input. We first evaluated *F(t)* across all the recorded terminals within one field of view at a given time *t* :

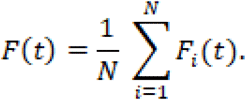

The variance of the time fluctuations of *F(t)* is

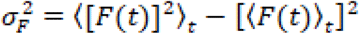

Where ⟨…⟩_*t*_ denotes the time-averaging over the session. For each terminal *F*_*i*_*(t)*, we used similar approach to calculate the time fluctuations

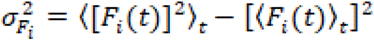

The synchrony measure, *χ*(*N*), for the scan filed is then calculated as

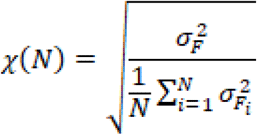

The value of *χ*(*N*) ranges between 0 and 1. *χ*(*N*) = 1 indicates that all ROIs within a scan are perfectly synchronized.

### Analysis of cone recordings

To control for possible effects of our pharmacological manipulation on outer retinal processing, we recorded calcium responses from the synaptic output pedicles of the four types of cones as described previously^7^ under control conditions and again following application of the GABA/glycine blocker cocktail, each time assessing cones’ responses to the spectral steps stimulus (Figure S4 C–N). For each cone, response amplitudes per step were extracted as described previously^7^, and tuning curves were computed by sign-inverting response amplitudes (to compensate for the hyperpolarising light response of vertebrate cones) and normalising each curve to a peak of 1.

## QUANTIFICATION AND STATISTICAL ANALYSIS

### Statistics

No statistical methods were used to predetermine sample size. Owing to the exploratory nature of our study, we did not use randomization or blinding.

Chi-squared tests were used for the following datasets: polarity and kinetics based on white step stimulation for ACs (Figure 1G); BCs polarity distribution before and after disinhibition with white steps (Figure 5A,B); BC response types before and after disinhibition (Figure 5E,F, bar plots); BCs polarity distribution before and after disinhibition with colour steps (Supplemental Figure S5); Polarity distribution for paired BCs (Figure 7E); paired BCs polarity distribution before and after sham injection with white steps (Figure S7 Extended H).

Wilcoxon signed-rank tests were used for the following datasets: comparison of population synchronicity (Supplemental Figure S4B); weight distributions for the four kinetic components (Figure 5D); population synchronicity in sham dataset (Supplemental Figure S7 Extended I); co-variation of absolute response amplitudes changes (Figure 8F,G, Supplemental Figure S8A-F).

Wilcoxon rank sum tests were used for the following datasets: distribution of kinetic component weights for ACs (Figure 1H); drug effect for four types of cones (Supplemental Figure S4E,H,K,N); Distribution of kinetic component weights for BCs (Figure 5E,F, distributions); Percentages of ROIs per spectral category (Figure 6B).

Two sample Kolmogorov–Smirnov tests were used for the following datasets: Distribution of On and Off BC-terminals across the IPL based on white steps (Figure 5C, D); Distribution of opponent terminals across the IPL based on spectral kernels (Figure 6C, D).

## SUPPLEMENTAL VIDEOS

**Supplemental Video 1 – related to Figure 4**. Example average-response movie of a central (left) and peripheral scan region (right) during white chirp stimulation (cf. Figure 4C,F)., in the presence (bottom) and absence (top) of inhibitory inputs from amacrine cells. To highlight increases SyjGCaMP8m signal relative to baseline, the movie is centered around each pixel’s mean brightness immediately preceding the 100% white step. 4x Real time.

**Supplemental Video 2 – related to Figure 4**. As Supplemental Video 1, but for colour-steps (cf. Figure 4D,G). The marker in the center indicates the current stimulus played. 4x Real time. To highlight both increases and decreases in the SyjGCaMP8m signal, the movie is centered around each pixel’s mean value across trials.

---

**Supplemental Figure S1 – related to Figure 1.**
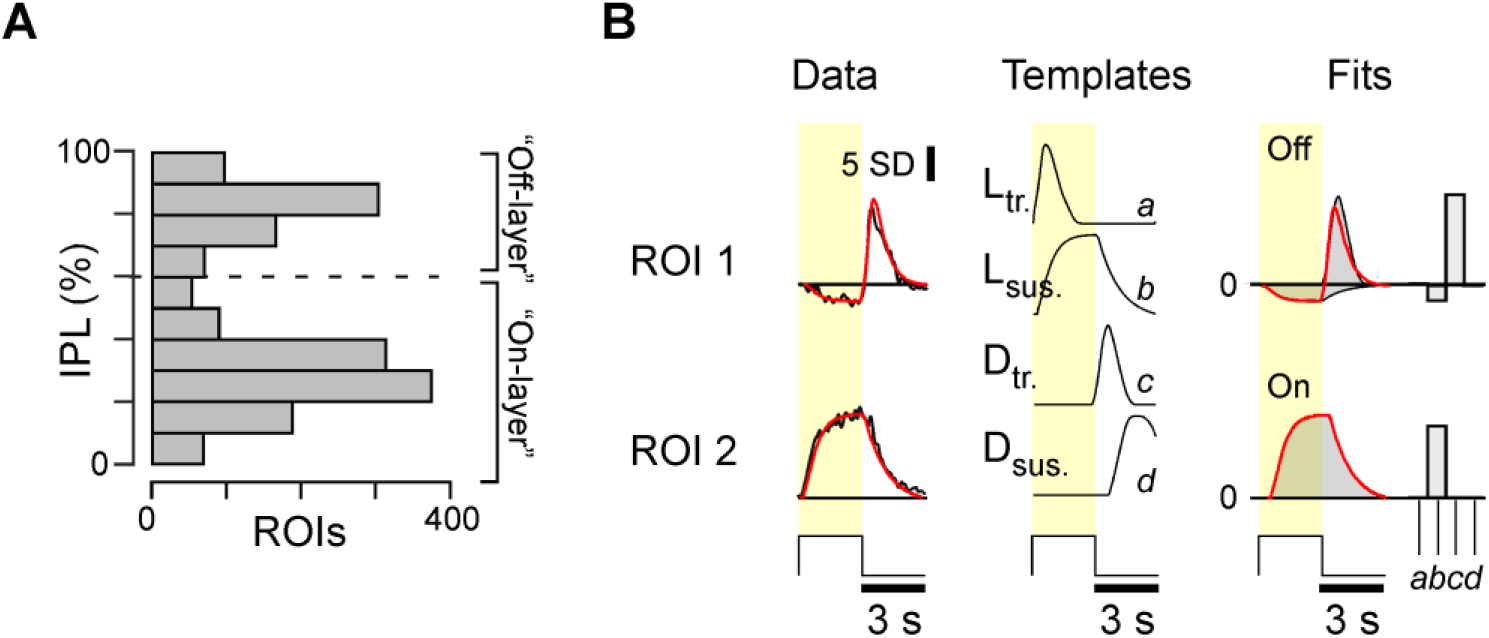
**A**, distribution of all AC-ROIs across the IPL. **B**, Kinetic template-fitting approach (Methods) illustrated using the white-step responses of example ROIs 1 and 2 (Figure 1D). Left: Response means (black) shown with the fit superimposed (red). Middle: Four kinetic templates are used for fitting: Light-transient, Light-sustained, Dark-transient, and Dark-sustained. Right: scaled kinetic components used to fit the two example ROIs (grey) with their sum superimposed (red), and corresponding component weights (bars).

**Supplemental Figure S2 – related to Figure 2.**
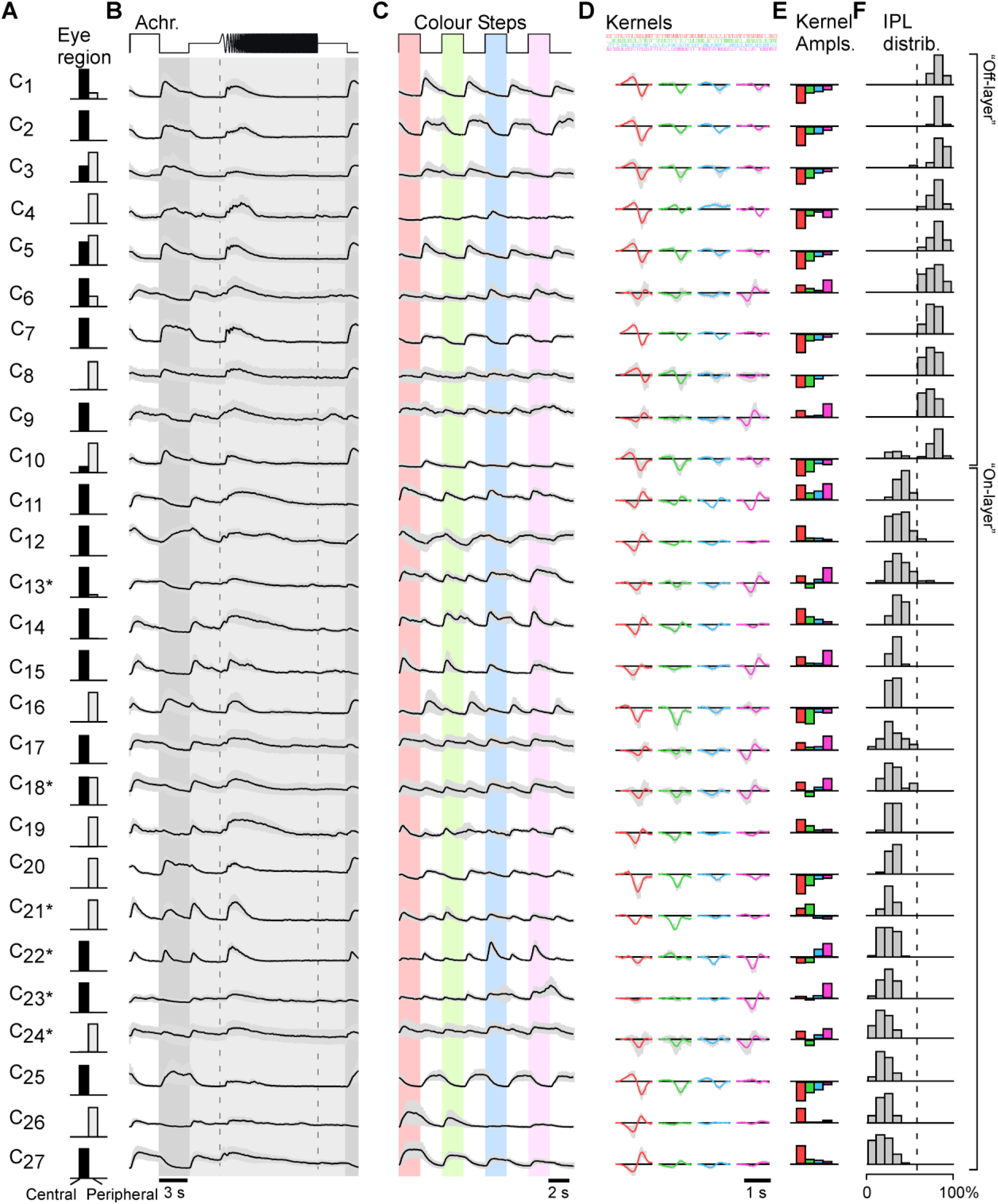
Full cluster overview of ACs. Shown are the eye region of the included ROIs (A, Central / Peripheral), as well as the mean±SD of chirps (B), colour-steps (C), Kernels (D), mean kernel amplitudes (E) and IPL distribution of included ROIs normalised to the population of all recorded ROIs (cf. Supplemental Figure S1A).

**Supplemental Figure S3 – related to Figure 3.**
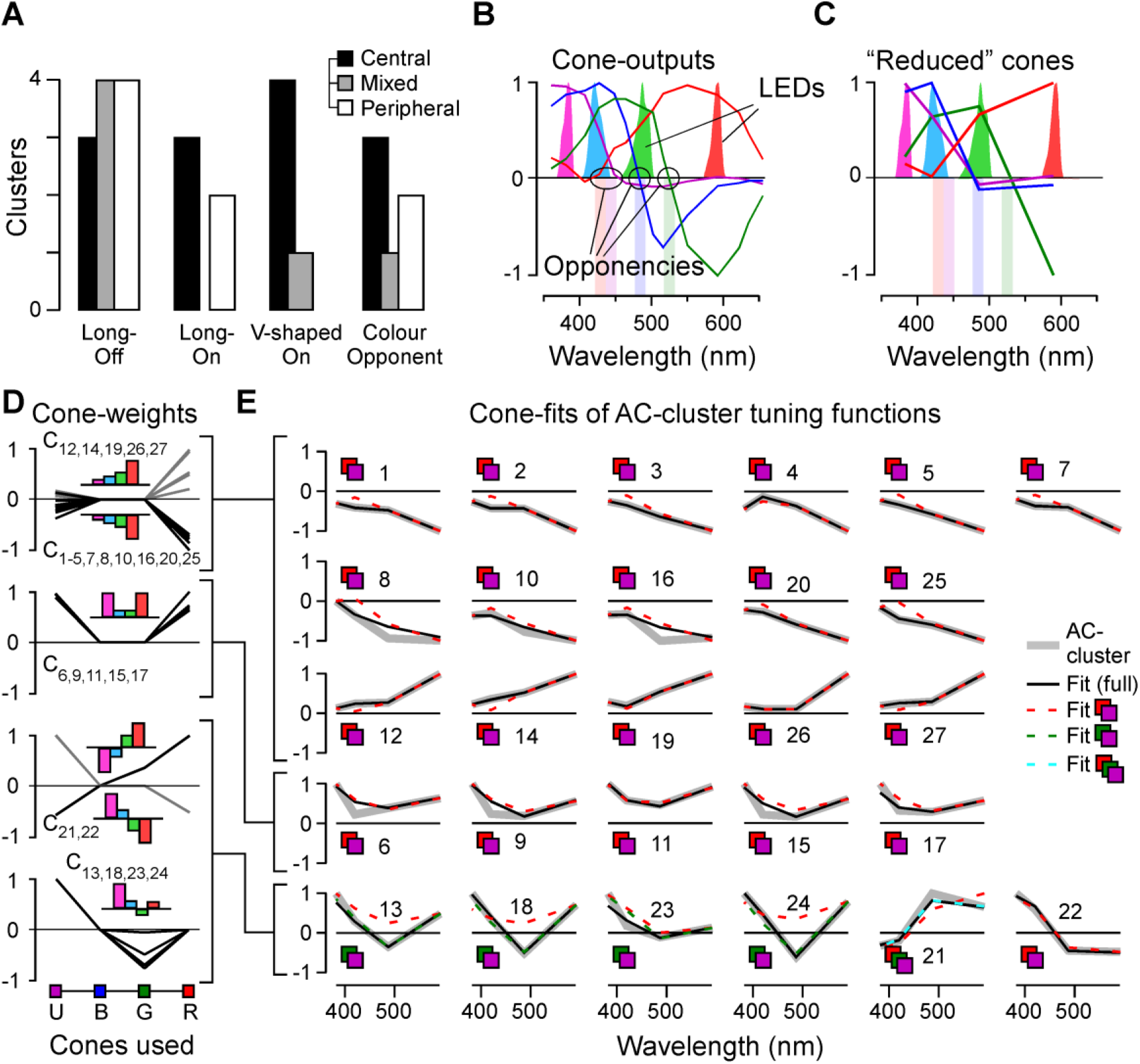
**A**, Distribution of AC-clusters across the four spectral groups (cf. Figure 3A-D), divided by retinal location: Central (black), Peripheral (white) and mixed (grey). **B**, Spectral relationship of cone-photoreceptor outputs (thin lines, based on Ref^7^), their spectral zero crossings (shaded bars, labelled “opponencies”), and the spectra of the four LEDs used to probe ACs and BCs in the present study (solid curves). Note that the spectral zero crossings of blue-cones coincides with the bulk of the spectral power of the green-LED. **C**, Reduced spectral tuning functions of the cone outputs as they would appear if probed with the four LEDs used in the present study. Note that the specific LED placement resulted in an under sampling of the full blue-cone opponency. **D**,**E**, Cone-weights (D) and overview of fits (E) between spectral tuning functions of reduced cones (see above) and AC-cluster means. The four plots in (D) part-correspond to the four spectral groups shown in Figure 3A-D, however with On- and Off versions of long-wavelength biased clusters combined (D, top), while the five colour-opponent clusters were further subdivided into two separate groups as shown (D, bottom). (E) shows one panel per cluster as indicated, sorted by spectral groups. Shown are: AC-cluster mean (grey, thick), the best fit when using all four cones (black) and the fit result when only using red- and UV-cones (red, dashed). We also show the fit results for the six opponent clusters (bottom row) when using only green- and UV-cones (green, dashed) and when using red-, green- and UV-cones (light blue, dashed). For quantitative evaluation of fits, see Methods.

**Supplemental Figure S4 – related to Figure 4.**
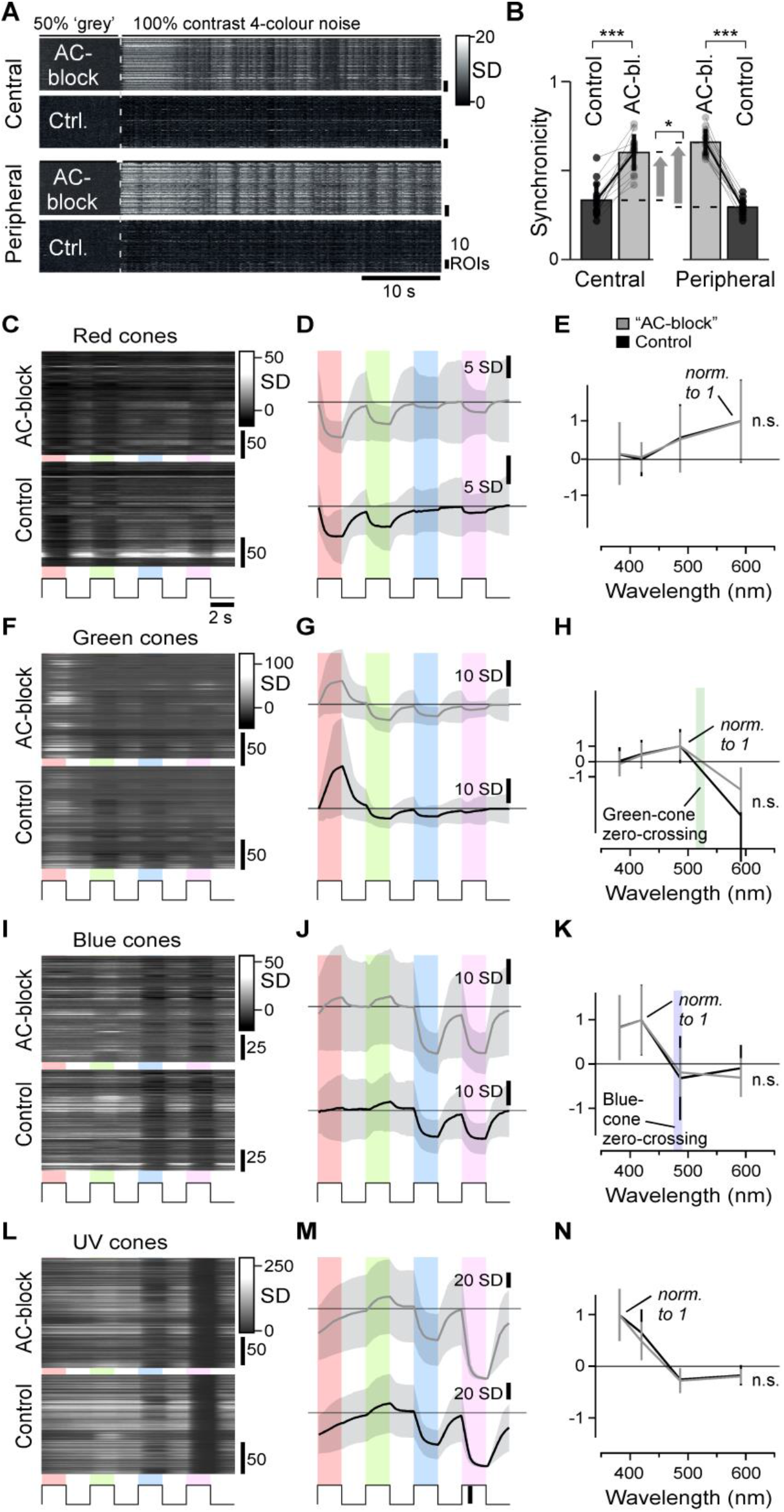
**A**,**B**, comparison of population synchronicity during spectral noise stimulation. Heatmaps show the first 60 s of the z-normalised responses example scans during control conditions and following AC-block, as indicated, for the central retina (top two rows) and the peripheral retina (bottom two rows). Population synchronicity (B) was computed as in Ref^63^. Wilcoxon Signed-Rank test: Central: p < 0.001; Peripheral: p < 0.001; Difference in change between central and peripheral: p = 0.017. **C-N**, No effect of drug-cocktail injection as in Figure 4 on spectral tuning including opponency in cones. Heatmaps (C) of SyGCaMP6f responses in red cone terminals to the four steps of light as Figure 4D based on previously established protocols^7^ before (bottom) and after drug injection (top), the same data summarised with mean±1SD shading (D) and extracted response amplitudes±1SD plotted against stimulus wavelength, with each curve’s peak hyperpolarising response normalised to 1 (E). **F-N**, as (C-E), but for green (F-H), blue- (I-K) and UV-cones (L-N), respectively. The coloured shadings in H and K indicates the spectral position of the green- and blue-cones’ zero-crossings, respectively, from Ref^7^. Wilcoxon rank sum tests were used for comparing amplitude-normalized tuning functions, p > 0.05 for all the four types.

**Supplemental Figure S5 – related to Figure 5.**
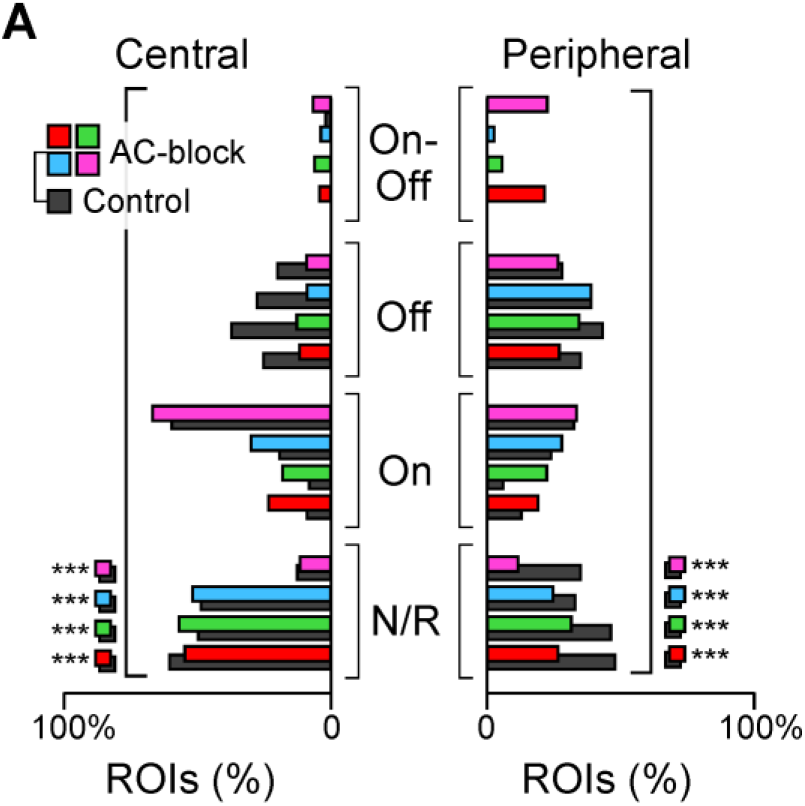
**A**, As bar plot in Figure 5A,B, but here shown separately for the four colour steps (cf. Figure 4D) instead of the single white step. Chi-squared tests were used to test for within-colour changes in the distribution of terminals by polarity, p < 0.001 for all four colours.

**Supplemental Figure S6 – related to Figure 6.**
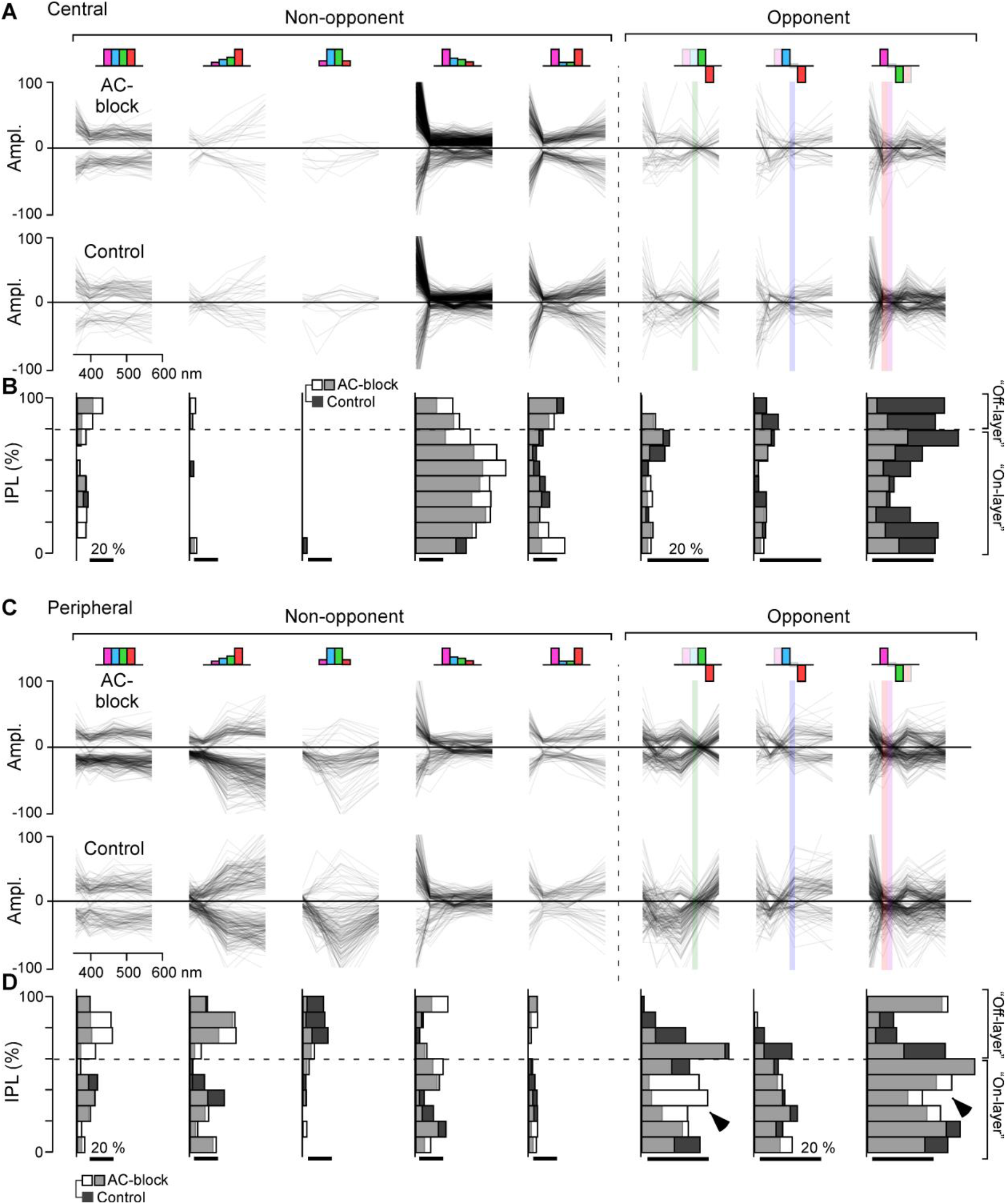
**A**, All central BCs’ spectral tuning functions (based on the kernels, cf. Figure 4E) under control (bottom) and AC-block condition (top) sorted into nine categories as indicated (Methods, cf. Figure 6A,B, plotted in same order). Individual BCs are plotted as semi-transparent grey such that darker shades overall indicate larger numbers of BCs. The coloured vertical bars in the three opponent groups (right) indicate their corresponding spectral zero crossings (based on Refs^7,18^ – cf. Supplemental Figure S3B,C). **B**, As Figure 6C,D, data from (A) summarised by IPL position. **C**,**D**, as (A,B), respectively, but for Peripheral retina. Arrowheads indicate IPL regions where the representation of spectral opponency systematically increases following AC-block. Two sample Kolmogorov–Smirnov tests, Central, p > 0.05; Peripheral, p < 0.001.

**Supplemental Figure S7 – related for Figure 7.**
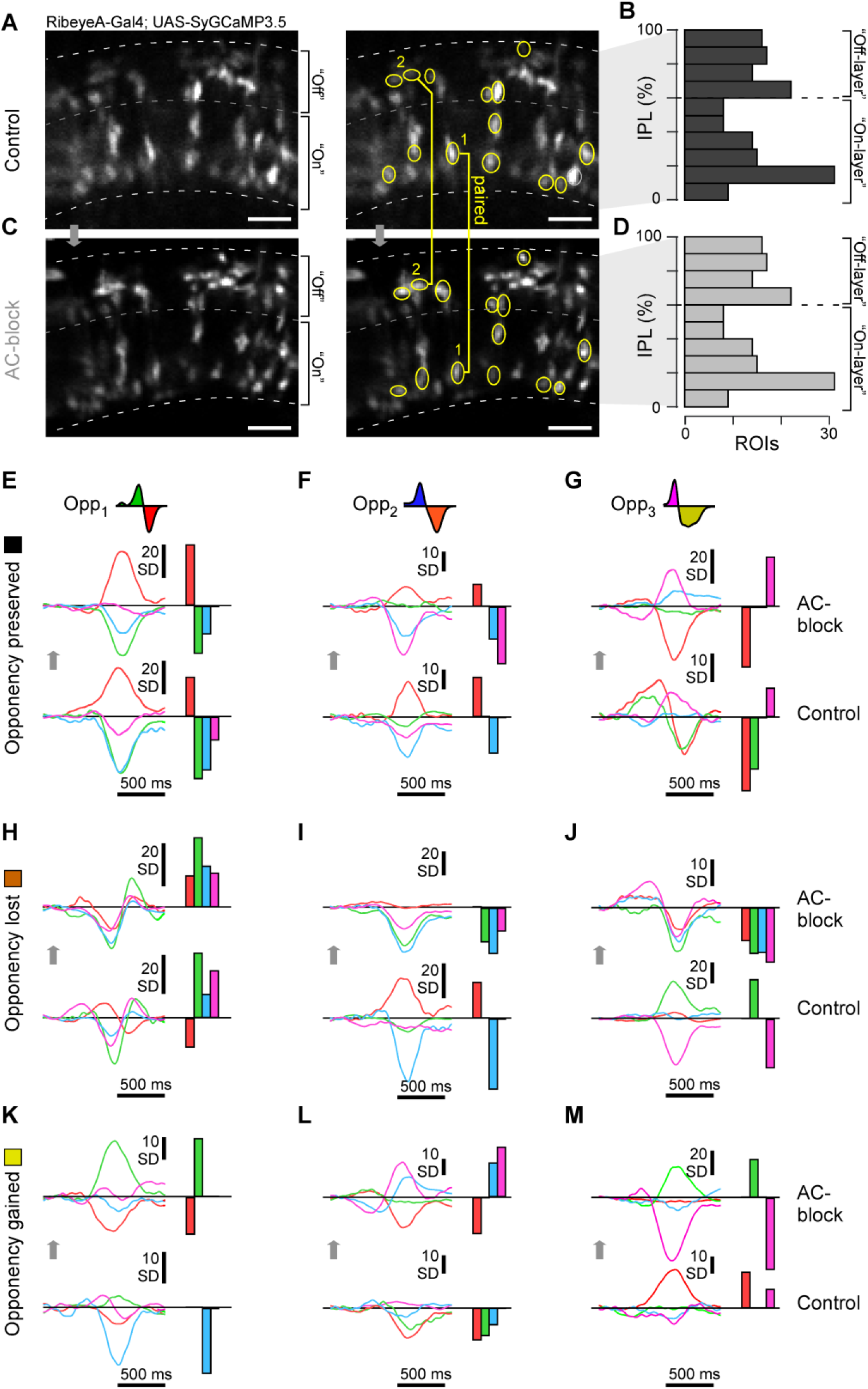
**A-D**, Example scan field with BCs sparsely expressing SyGCaMP3.5 under control conditions (A) and the same field of view following AC-block (B). Approximately half of visible BC terminals could be reliably matched across the two conditions (yellow) and were counted as paired data. Paired terminals spanned the entire depth of the IPL (B,D). **E-M**, Selected example BCs (paired data) that either preserved (E-G), lost (H-J) or gained (K-M) spectral opponency following pharmacological removal of inner retinal inhibition. Examples from all three types of opponencies are presented: red:green (Opp_1_, E,H,K), red/(green):blue (Opp_2_, F,I,L), (red)/green:UV (Opp_3_, G,J,M). Shown in each case are the four spectral kernels (left) and their automatically extracted response amplitudes (right, Methods).

**Supplemental Figure S8 – related to Figure 9.**
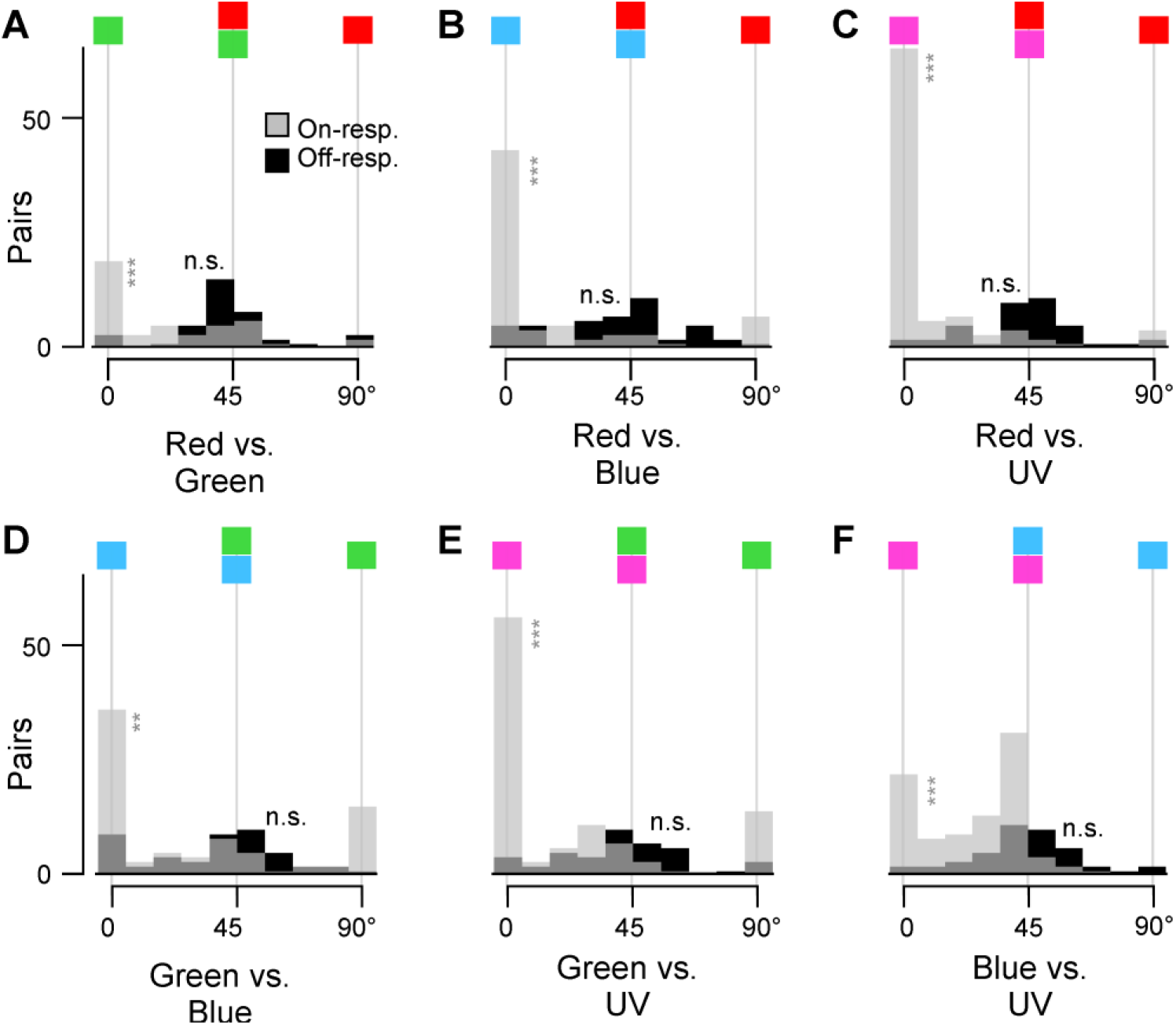
**A-F**, as Figure 9F, but for all six possible colour combinations. Wilcoxon Signed-Rank tests On: p < 0.001 for all the combinations; Off: p > 0.05 for all the combinations.

## EXTENDED DATA

**Supplemental Figure S2 extended 1 – related to Figure 2.**
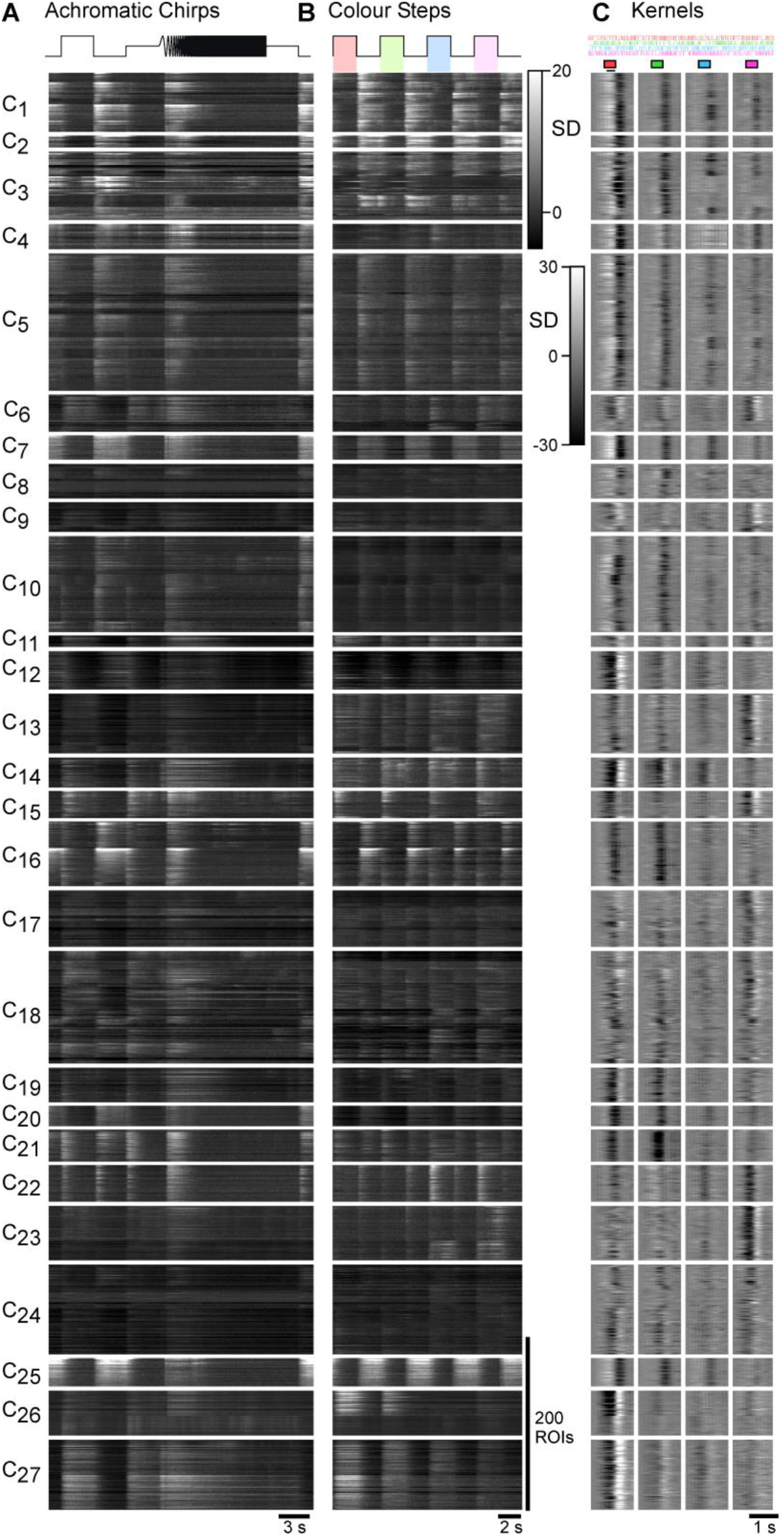
Detail of AC clusters, showing heatmaps of the full dataset leading to the cluster means shown in Supplemental Figure S2. Shown are the chirps (A), colour-steps (B), and kernels (C).

**Supplemental Figure S2 extended 2 – related to Figure 2.**
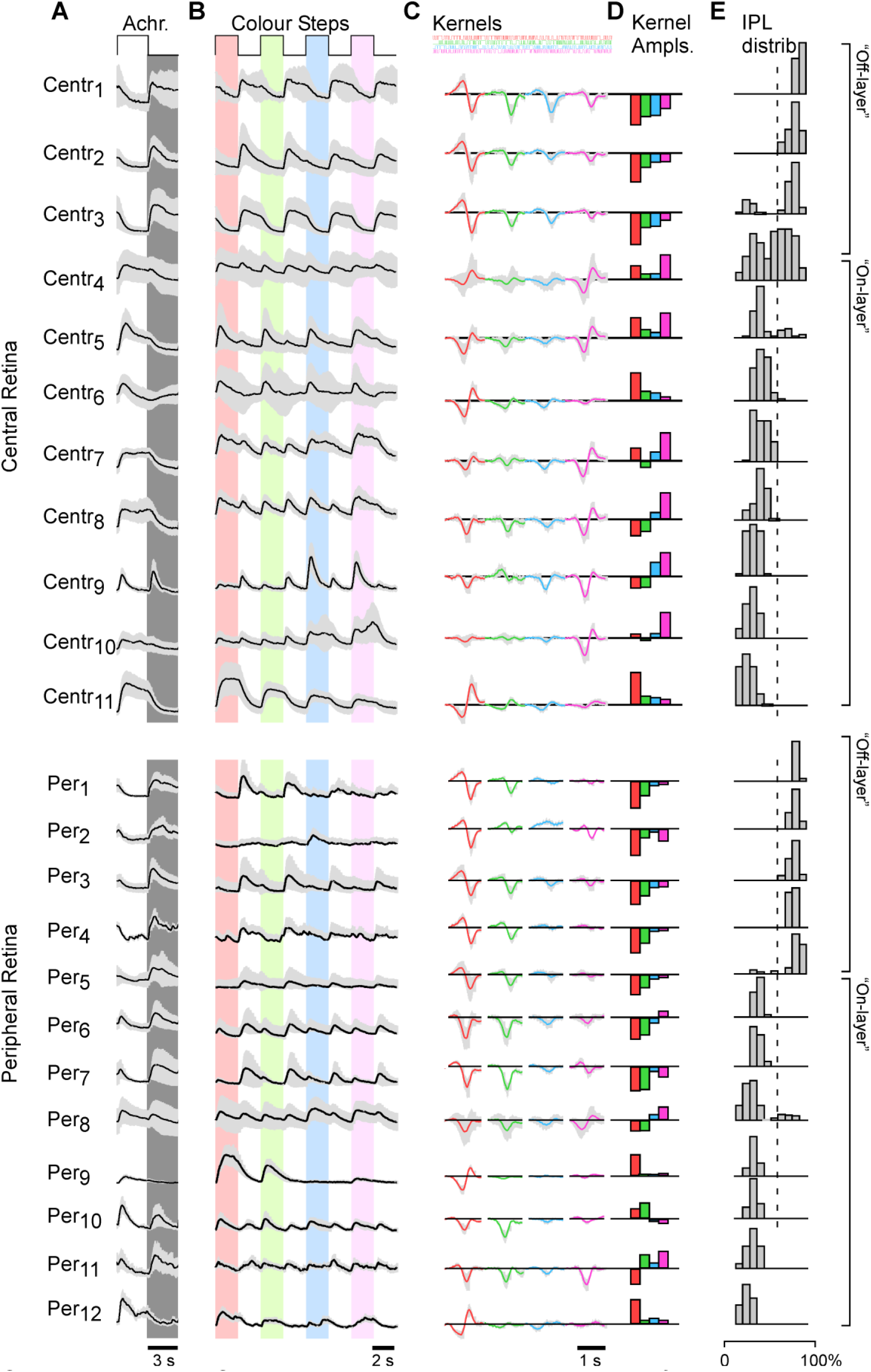
Result of alternative clustering, where ROIs from the central and peripheral retina were clustered independently.

**Supplemental Figure S7 Extended – related for Figure 7.**
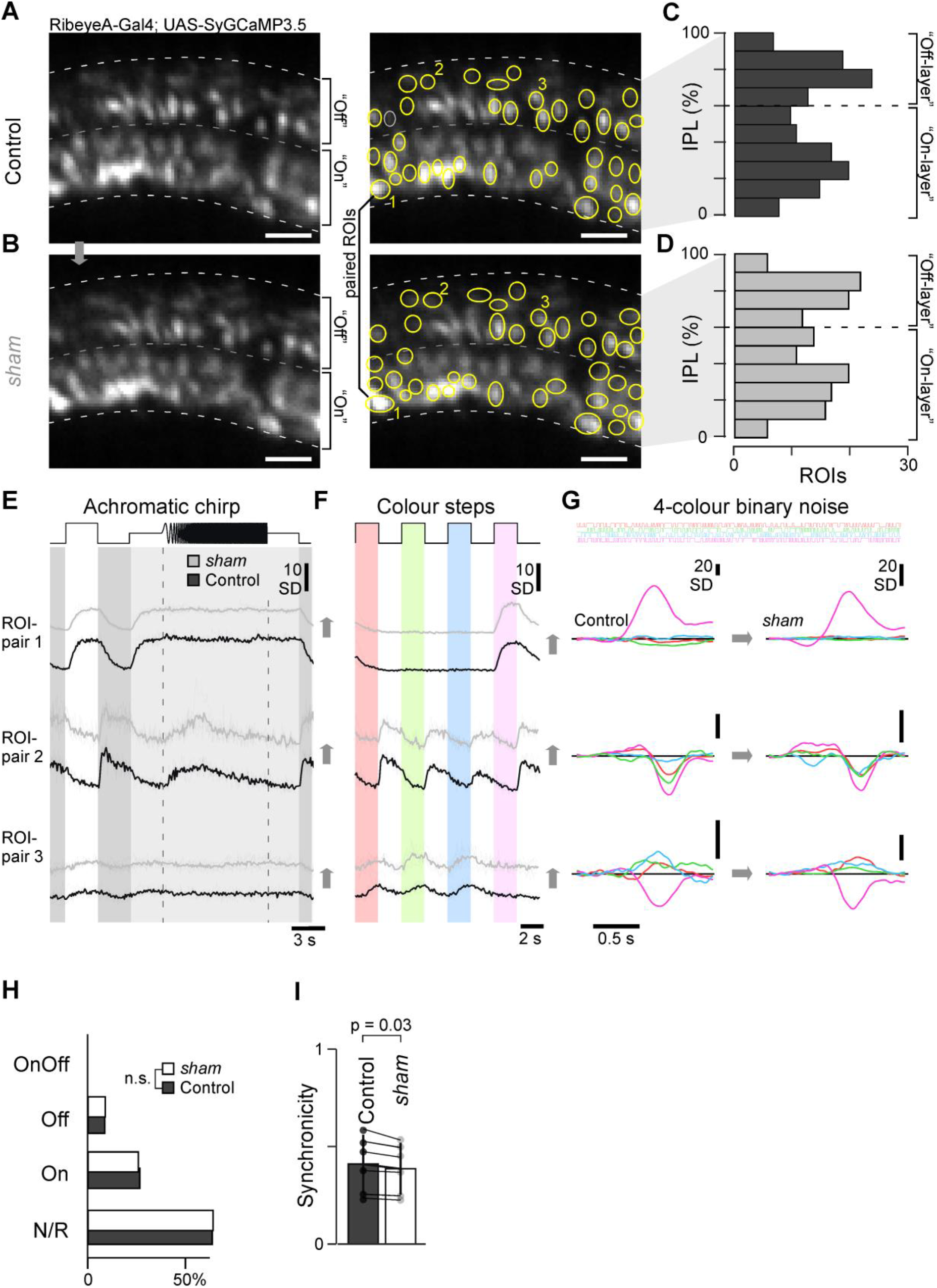
**A-G**, As Supplemental Figure S7A-D and Figure 7A-C, respectively, but here shown for sham injection dataset. **H**, No change in response allocation based on the white step responses in the sham dataset before (dark grey) and after sham injection (white). Chi-squared test, p > 0.05. **I**, No change in population synchronicity in sham dataset. Wilcoxon Signed-Rank test: p = 0.03. (cf. Supplemental Figure S4B).

## Acknowledgements

We thank Thomas Euler for critical feedback. The authors would also like to acknowledge support from the FENS-Kavli Network of Excellence and the EMBO YIP. Funding was provided by the Wellcome Trust (Investigator Award in Science 220277/Z20/Z to TB and 102905/Z/13/Z to LL), the European Research Council (ERC-StG “NeuroVisEco” 677687 to TB), UKRI (BBSRC, BB/R014817/1 to TB), the Leverhulme Trust (PLP-2017-005 and RPG-2021-026 to TB) and the Lister Institute for Preventive Medicine (to TB).

This research was funded in whole, or in part, by the Wellcome Trust [220277/Z20/Z and 102905/Z/13/Z to LL]. **For the purpose of Open Access, the authors have applied a CC BY public copyright licence to any Author Accepted Manuscript version arising from this submission**.

