## Supplementary figures and images for "Amacrine cells differentially balance zebrafish colour circuits in the central and peripheral retina"

### Supplemental Video 1

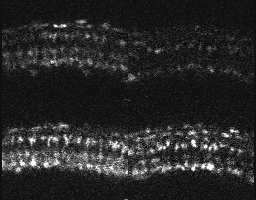

### Supplemental Video 2

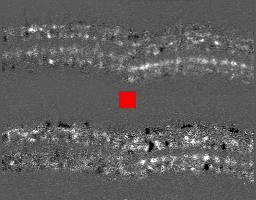
